# Nuclear Cx43 restrains microglial neurotoxicity during brain development

**DOI:** 10.64898/2026.06.29.735292

**Authors:** Yixun Su, Qiumin Feng, Qi Wang, Parsa Khakpour, Hui Li, Chenmeng Wang, Xiaoying Chen, Zhonghao Wu, Shaoqi Zhu, Marie-Eve Tremblay, Rao Fu, Hui Chen, Jianqin Niu, Alexei Verkhratsky, Chenju Yi

## Abstract

Microglia are essential for sculpting the developing brain, yet the molecular mechanisms that select beneficial overreactive phagocytosis remain incompletely understood. Connexin 43 (Cx43, encoded by *GJA1* in humans) is best known as a gap junction and hemichannel protein, although its non-canonical, channel-independent functions are increasingly recognized. We found that Cx43 is highly expressed in microglia during the perinatal period in human and mouse, whereas proportion of full-length multimeric Cx43 unexpectedly localizes to the nucleoplasm. Deletion of microglial Cx43 in mice during development instigates a transient neurotoxic state with microgliosis, upregulated phagocytic and complement pathways, excessive neuronal apoptosis, translating into depressive-like and cognitive deficits in the adulthood. Notably, neither microglia-specific deletion of Cx43 in adulthood nor hemichannel blockade recapitulate these changes, indicating a channel-independent, developmental stage-specific neuroprotective mechanism. Nucleus-targeted Cx43 overexpression suppresses neurotoxic markers and neural apoptosis. Nuclear Cx43 interacts with transcriptional regulators to restrain proinflammatory gene programs, nuclear import of Cx43 is driven by neurogenic niche-derived bFGF, which triggers AKT-mediated phosphorylation of a C-terminal nucleus localization signal (NLS), 14-3-3 binding, and importin-dependent nucleus translocation. These findings reveal a developmentally restricted nuclear Cx43 function that restrains microglial neurotoxicity while promoting microglial physiological functions thus expanding connexin biology to transcriptional co-regulation and pointing to a potential avenue for therapeutic intervention.

## INTRODUCTION

Microglia, the resident immune cells of the central nervous system (CNS), colonize the brain during early embryonic development^1^ ^2^, and, beyond immune surveillance, play critical roles in shaping the developing brain by removing excessive neurons and oligodendrocyte precursor cells ^3^ ^4^, pruning synapses ^5^, and remodeling myelin ^6^. These functions are tightly regulated by the local environmental cues, particularly secreted factors from neural progenitor cells including vascular endothelial growth factor (VEGF) and basic fibroblast growth factor (bFGF) that influence microglial proliferation, morphology, and phagocytosis ^7^. Disruption of these functions during development translate into neurodevelopmental and neuropsychiatric disorders ^6^ ^8^ ^9^. Microglial phagocytosis in the healthy developing brain is largely beneficial and protective; however, when overactivated, as seen in neurodegeneration in response to abnormal neurotoxic stimulus, it can impair neuronal survival and neurite outgrowth through proinflammatory pathways ^10^ ^11^. The developing CNS thus likely employs dedicated mechanisms to constrain maladaptive phenotypes. For example, the transcription factor Myocyte-specific enhancer factor 2C (MEF2C) prevents neurotoxic microgliosis during normal development ^12^, yet the full repertoire of molecular brakes that maintain microglial homeostasis in the perinatal brain remains incompletely understood.

Connexin 43 (Cx43, encoded by *GJA1* in humans and *Gja1* in mice) is a transmembrane protein best known for forming gap junction channels and hemichannels at the plasma membrane. Although Cx43 is predominantly studied in astrocytes ^13^, it is also present in microglia, where it operates as a hemichannel to facilitate the release of signaling molecules, such as ATP, and contribute to neuronal damage under pathological conditions ^14^ ^15^; of note microglial Cx43 do not form gap junctions *in vivo* ^16^. Recently, Cx43 has been recognized to possess non-canonical functions linked with its subcellular localization. For example, membrane Cx43 could serve as an adhesion molecule ^17^ or act as a scaffolding platform via its C-terminus ^18^ ^19^; Cx43 mitochondrial translocation could modulate oxidative phosphorylation ^20^ ^21^; Cx43 can be released in extracellular vesicles and promote exosome uptake by target cells ^22^. Truncated isoforms of Cx43 can translocate to the nucleus ^23^ ^24^, while full-length Cx43 were also detected at the nuclear envelope of cardiomyocytes ^25^ ^26^, where they regulate gene transcription. Whether microglial Cx43 plays a role in brain development through its non-canonical functions remains unknown.

In this study, we examined the profile of microglial Cx43 in human and mouse developing brains and identified a previously unrecognized nuclear pool of full length, multimeric Cx43 that peaks during perinatal development. We found that nuclear Cx43 actively monitors the neurotoxic microglial gene program through its interaction with key transcriptional regulators. Conditional deletion of microglial *Gja1* during embryonic or postnatal development, but not in adulthood, leads to excessive neuronal apoptosis, depletes the hippocampal neurogenic pool, resulting in life-long depressive-like behaviors and cognitive deficits. Mechanistically, we show that neurogenic niche derived signals, such as bFGF drive AKT-mediated phosphorylation of the C terminus of nuclear localization signal (NLS), promoting 14-3-3 binding and importin-dependent nuclear import of Cx43. These findings identify a developmentally restricted nuclear function of Cx43 that acts as a molecular brake on microglial neurotoxicity and establish connexin-dependent transcriptional coregulators in the developing brain.

## RESULTS

### Profiling of microglial Cx43 expression and function in developmental brain

First, we examined the expression of Cx43 in human microglia during development. We analyzed a dataset with > 700,000 snRNA-seq profiles sourced from 169 brain samples and 106 donors, with developmental stages ranging from the 2^nd^ trimester of pregnancy to the adulthood ^27^ (Fig. 1A). In this dataset, we found low microglial Cx43 (*GJA1*) levels at 4^th^ Last Menstrual Period (LMP), which were gradually increased during embryonic development, peaked at birth, and then gradually decreased during postnatal development (Fig. 1B). We also examined the expression of other connexin family genes, including Cx36 (*GJD2*) and Cx32 (*GJB1*), reported in microglia ^28^ ^29^, as well as Cx26 (*GJB2*) and Cx30 (*GJB6*) enriched in astrocytes ^30^. In microglia, compared to Cx43, expression of all these genes was lower, and it did not change during development (Fig. 1B). To verify microglial Cx43 expression during development, we performed immunostaining on a cryosection of human embryonic brain tissue (gestation week 19). It was observed that at this stage, microglia were enriched in the subventricular zone (SVZ) and dispersed into the intermediate zone (IZ) and subplate zone (SP), but were largely absent from the cortical plate (Fig. 1C). We confirmed the colocalization of Cx43 signal within IBA1+ microglial domain; there was no significant difference in the amount of Cx43 in microglia from SVZ, IZ and SP (Fig. 1C, D, Fig. S1A).

**Fig. 1.**
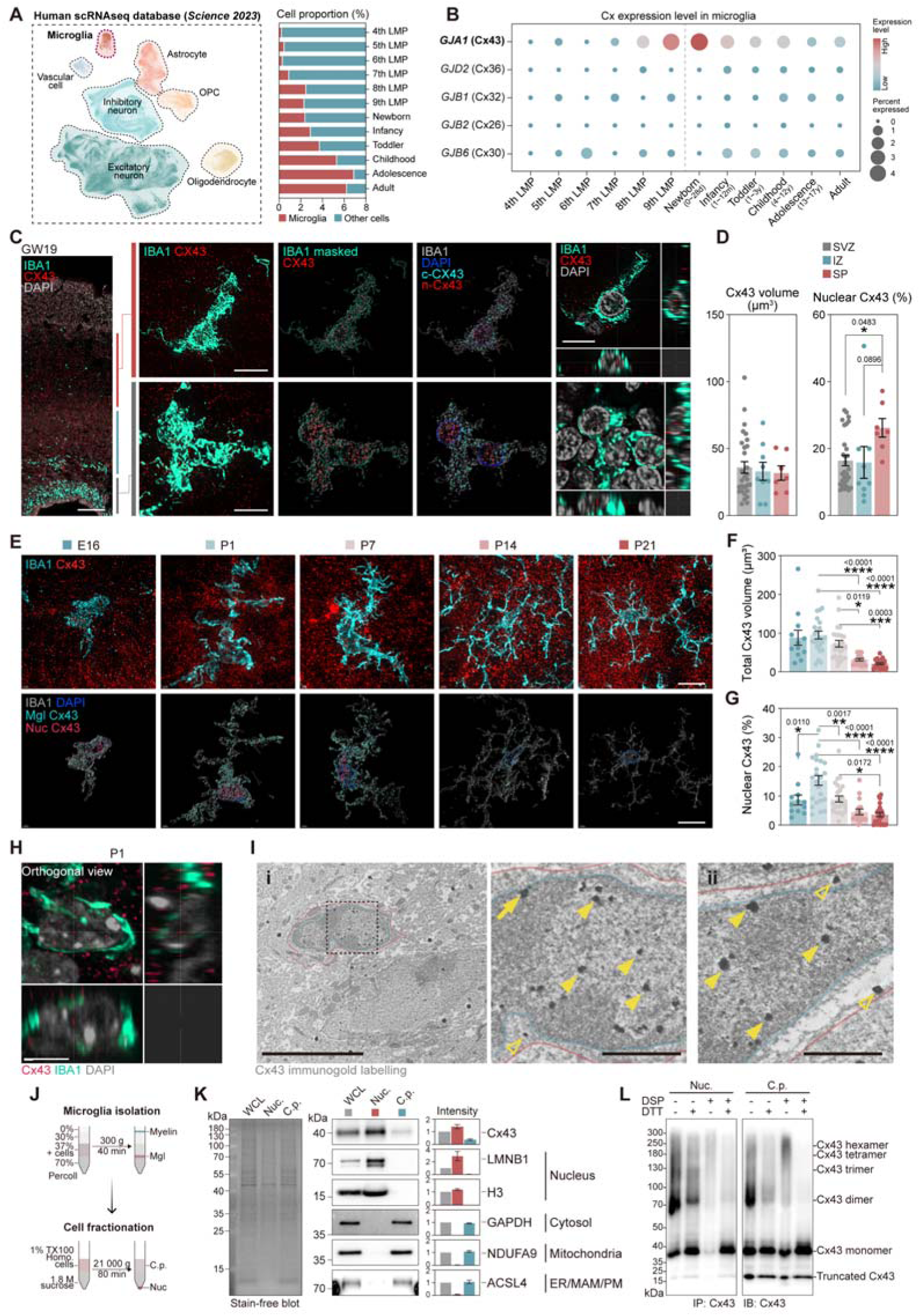
Developmental profiling of microglial Cx43 expression and function. **(A)** snRNA-seq analysis during human brain development ^27^. UMAP visualization and cell type annotation, and percentage of microglia in sample from different stages. LMP, last menstrual period. **(B)** Dot plot of Cx encoded gene expression in microglia at different stages. Dot color indicates the expression level, and the size shows the percentage of cells expressing the gene. **(C)** Representative images of IBA1 and Cx43 fluorescence staining in human embryonic cortex at gestation week 19 (GW19) (Scale bar, 500 μm). Two representative super-resolution images of IBA1^+^ microglia from subventricular zone (SVZ) and subplate zone (SP) were shown, together with the 3D reconstructions of IBA1/DAPI masked Cx43 signal, and the orthogonal views (Scale bar, 10 μm). **(D)** Quantification of IBA1-masked Cx43 volume and the percentage of nucleus Cx43, in SVZ, intermediate zone (IZ), and SP. N = 29, 9, and 7 cells. **(E)** Representative fluorescence super-resolution images of mouse cortex at different stages (E16, P1, P7, P14, and P21), staining for Cx43 and IBA1, together with the 3D reconstructions of IBA1/DAPI masked Cx43 signal (Scale bar, 10 μm). **(F)** Quantification of IBA1-masked Cx43 volume. **(G)** Quantification of the percentage of nucleus Cx43. N = 12, 23, 25, 17, and 21 cells. **(H)** Orthogonal view of the P1 microglia in (E) (Scale bar, 5 μm). **(I)** Representative immunoelectron microscopy images of two microglia (**i** and **ii**) in dorsal hippocampus, *stratum lacunosum moleculare* from a P7 wild type mice, immunogold-labeling for Cx43. Red line, microglia cell body. Green line, microglia nucleus. Hollow arrowhead, Cx43 signal in the cytoplasm. Arrowhead, Cx43 in the nucleus. Arrow, Cx43 at the nuclear envelope (Scale bar, 5 μm and 1μm). **(J)** Schematic of microglia isolation and fractionation. (K) Western blot analysis of microglia fractionations, blotting for Cx43, nucleus marker (LMNB1, H3), Cytosol marker (GAPDH), Mitochondria marker (NDUFA9), and membrane marker (ACSL4). Stain-free blot was shown. WCL, whole cell lysate. Nuc., nucleus. C.p., cytoplasm. N = 3 experiments. **(L)** Western blot analysis of Cx43-immunoprecipitated microglia nucleus and cytoplasmic fraction, with or without DSP crosslinking or DTT decrosslinking, blotting for Cx43. Cx43 multimers were labeled at their predicted molecular weight. Results are expressed as mean ± SEM. Significance: \**p* < 0.05, \*\**p* < 0.01, \*\*\**p* < 0.001, \*\*\*\**p* < 0.0001.

Next, by examining the mouse microglial Cx43 expression in a single-cell RNA-seq dataset ^31^, we found that Cx43 was highly expressed in microglia at both the embryonic and perinatal stages; Cx43 expression declined in adulthood (Fig. S1B). The high Cx43 mRNA expression during early postnatal (P) stages (P0∼P7) was confirmed using RT-qPCR from acutely isolated microglia (Fig. S1C). To further verify microglial Cx43 protein, we immuno co-stained Cx43 with microglial marker IBA1 in brain sections from mice at embryonic day 16 (E16), P1, P7, P14, and P21. Analysis of Cx43 signal within IBA1+ domain showed a high Cx43 expression in developing microglia from E16 to P7, with an increase from E16 to P1, before decreasing and dropping to a low level after P14 (Fig. 1E, F). Cx43 expression negatively correlated with microglia ramification (Fig. S1D). Together, these data reveal a high Cx43 expression level in human and mouse microglia during perinatal development.

When analyzing microglial images, we noticed a non-canonical, nucleus localization of microglial Cx43 in perinatal development. In the human embryonic cortical region, abundant Cx43 signals (SVZ: 16.4 ± 1.5%, IZ: 15.8 ± 4.7%, SP: 26.2 ± 2.8%) were observed in microglial nuclei; this signal was higher in the SP compared to the SVZ (Fig. 1C-D). Similarly, Cx43 was also found in the microglial nuclei of developing mouse brains (Fig. 1E, G). The fraction of nuclear Cx43 was high at early postnatal days gradually decreasing as microglia mature (Fig. 1G). The Cx43 signals were localized to the nucleoplasm, but not to the nuclear envelope, in contrast to the previous finding in primary cardiomyocytes ^25^ (Fig. 1C, E, H). We further examined the Cx43 nucleus translocation using immunoelectron microscopy, which identified Cx43 in the nucleoplasm of microglia, particularly in the euchromatic region (Fig. 1I, Fig. S1E).

To better evaluate the amount of nucleus-localized microglial Cx43, we fractionized acutely isolated microglia to obtain the pure nucleus and the cytoplasm fractions ^25^, followed by western blot analysis with the same protein loading amount of each fraction (Fig. 1J). The full-length Cx43 was enriched in the nucleus fraction compared to the whole cell and the cytoplasm fraction (Fig. 1K). Truncated versions of Cx43 derived from alternative translation initiation have been shown previously to translocate to the nucleus ^23^ ^24^. Indeed, two weak bands of truncated Cx43 at ∼32 kDa and ∼20 kDa were also found in the nucleus, but their level was much lower compared to the full-length form (2.13 ± 0.82% and 7.48 ± 2.06% of full-length) (Fig. S1F). Thus, the full-length, but not truncated, Cx43 is present in the nucleus, in particular in the nucleoplasm, in microglia of developing brains.

Translocation of Cx43 into the nucleus was more prominent in microglia compared to other brain cell types. Fractionation of the whole brain cells also showed nuclear localization of Cx43, but at a lower level compared to microglia (Fig. S1G). We further compared the nuclear Cx43 level in microglia and astrocytes, in which Cx43 is prominently expressed, in primary mixed glial cultures. Immunocytochemistry showed that CD11b+ microglia exhibited 2.49 ± 0.63-fold higher nucleus Cx43 level compared to GFAP+ astrocytes (Fig. S1H).

Cx43 hexamers are assembled in the Golgi apparatus before being trafficked to the plasma membrane ^32^ ^33^. Cx43 residing in the nuclear envelop was shown to derive from the endocytosed surface Cx43 ^25^. To resolve the potential multimeric or monomeric state of nucleus Cx43, we performed a crosslinking assay with DSP followed by Cx43 pulldown and Western blot. Nuclear Cx43 in microglia predominantly exist in multimeric states, with only little proportion of monomeric Cx43 (See + DSP group) (Fig. 1L). The naïve, non-crosslinked condition also revealed a dimerized band of Cx43 (Fig. 1L). In contrast, Cx43 in the cytoplasm exists in both preassembled monomeric state and multimeric state (Fig. 1L). Thus, before translocating to the nucleus Cx43 was assembled into multimers at the Golgi apparatus.

### Developmental ablation of microglial Cx43 drives a neurotoxic microglia phenotype

To reveal the functional relevance of microglial Cx43 during development, we crossed *Cx3cr1^CreERT^* mice with *Cx43^flox^*mice to obtain the *Cx3cr1^CreERT/+^::Cx43^f/f^* mice (Cx43^mglcKO^), achieving microglia-specific Cx43 ablation under tamoxifen-induced Cre recombination. In light of the potential background effect of *Cx3cr1^CreERT^* strain ^34^ ^35^, the *Cx3cr1^CreERT/+^::Cx43^+/+^* and *Cx3cr1^CreERT/+^::Cx43^f/+^* littermates were used as controls. Tamoxifen was given from P3 to P5, and the brain histology were analyzed at P7 (Fig. 2A). Ablation of microglial Cx43 caused significant increase of microglia number in most of the brain regions, including hippocampus, motor cortex, lateral hypothalamus, and nucleus accumbens (Fig. 2B-C). In cortical regions of Cx43^mglcKO^ mice, we found a significantly increased number of microglia lining the layer I region (Fig. 2B). Morphological examination using Sholl analysis revealed a profound morphological change in Cx43-ablated microglia, which presented more ramified morphology (mean intersection number), in hippocampus, prefrontal cortex, motor cortex, amygdala, and nucleus accumbens (Fig. 2D, Fig. S2A). The morphological change of microglia under Cx43 ablation was also exemplified by Principal Component Analysis (PCA) of morphological trait derived from Sholl analysis (Fig. S2B). We also noticed that Cx43-ablated microglia display significantly increased number of phagocytic pouch (Fig. 2E-F), likely indicating increased phagocytic activity.

**Fig. 2.**
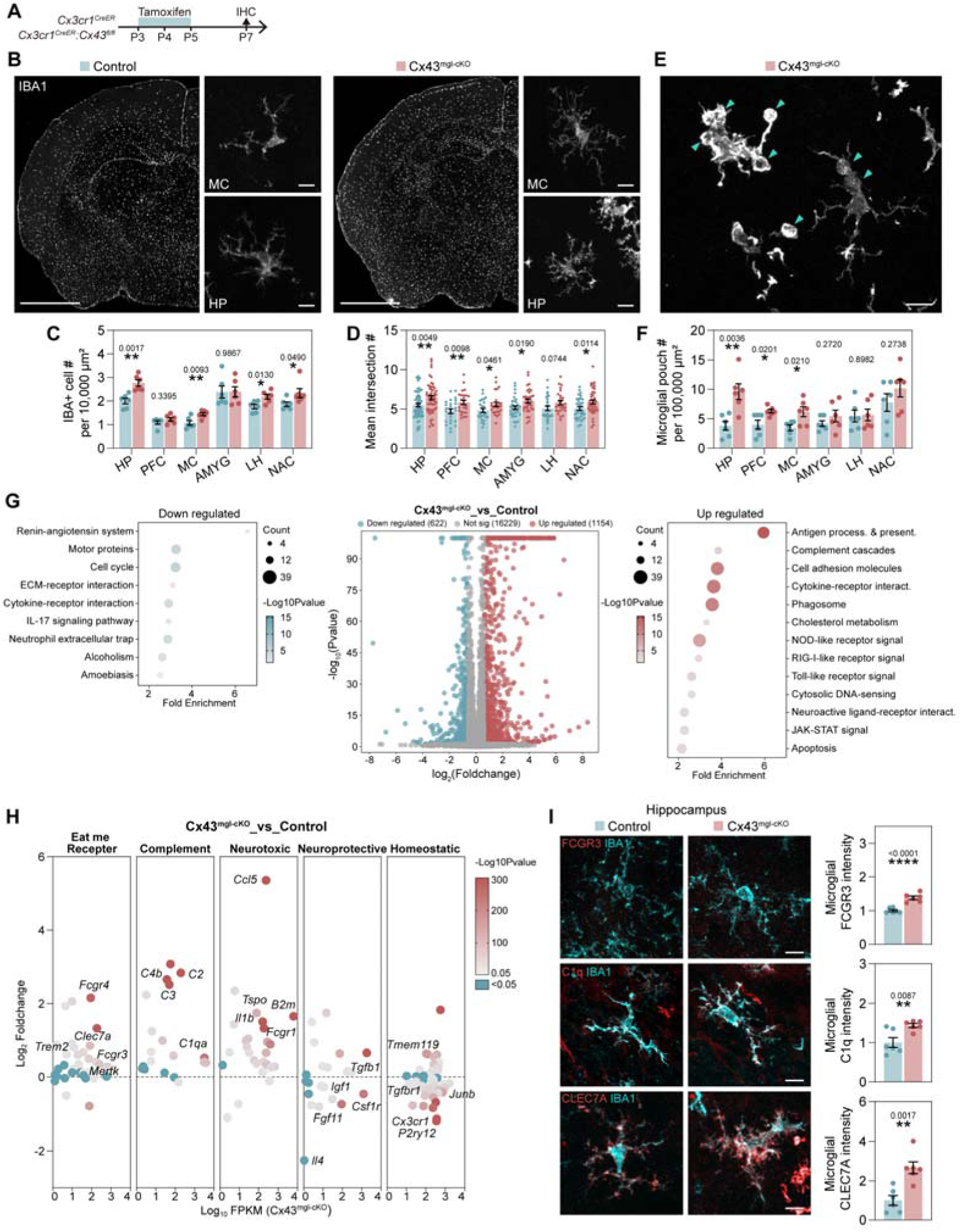
Early postnatal Cx43 ablation triggers a neurotoxic microglia phenotype. **(A)** Schematic of microglial Cx43 knockout induction and examination. **(B)** Representative images of IBA1 fluorescence staining in motor cortex (MC) and hippocampus (HP) from control and Cx43^mgl-cKO^ mice at P7. (Scale bar, 500 μm and 10 μm). **(C)** Quantification of IBA1^+^ cells in different brain regions. PFC, prefrontal cortex. AMYG, amygdala. LH, lateral hypothalamus. NAC, nucleus accumbens. N = 6 mice. **(D)** Quantification of mean intersection number from Sholl analysis of IBA1^+^ cells in different brain regions, N = 60, 55 (HP), 25, 25 (PFC), 25, 26 (MC), 32, 34 (AMYG), 28, 26 (LH), 35, 39 (NAC) cells from 6 mice. **(E)** Representative images of IBA1 fluorescence staining in the cortex from Cx43^mgl-cKO^ mouse at P7 (Scale bar, 10 μm). Arrowhead highlights the phagocytic pouch. **(F)** Quantification of microglial phagocytic pouch number, N = 6 mice**. (G)** RNA-seq analysis of acutely isolated microglia from the cortex and hippocampus of P7 control and Cx43^mgl-cKO^ mice. Dot plots show the KEGG pathway functional annotation of down-regulated and up-regulated genes. Volcano plot highlights the differentially expressed genes. N = 3 mice. **(H)** Change of genes involved in selective pathways. The color shows the p-value. **(I)** Representative images and quantification of IBA1, eat-me receptors (FCGR3 Clec7a), and C1q fluorescence staining in the hippocampus of P7 control and Cx43^mgl-cKO^ mice (Scale bar, 10 μm), N = 6 mice. Results are expressed as mean ± SEM. * *p* < 0.05, \*\**p* < 0.01, \*\*\*\**p* < 0.0001.

To further characterize microglial phenotype in Cx43^mglcKO^ mice, we acutely isolated microglia from the cortex and hippocampus of P7 control and Cx43^mglcKO^ mice, followed by RNA-seq analysis (Fig. 2G). Microglial Cx43 ablation caused profound transcriptomic changes in microglia, with upregulation of genes enriched in pathways such as antigen process & presentation, complement cascade, phagosome, apoptosis, etc. (Fig. 2G). To further characterize transcriptomic alterations of Cx43 knockout microglia, we surveyed the expression of genes involved in ‘eat-me’ signal reception, complement cascade, neurotoxicity, neuroprotection, and homeostatic genes. Microglia in Cx43 conditional knockout mice displayed upregulated genes involved in ‘eat-me’ signals, complement cascade, neurotoxicity, and downregulated genes associated with neuroprotection and homeostasis (Fig. 2H). Using immunostaining, we verified the upregulation of the ‘eat-me’ receptor FCFR3 and Clec7a, as well as the complement cascade component C1q in Cx43-ablated microglia using immunostaining, which aligned with the increased number of microglial phagocytic pouches (Fig. 2F, I, Fig S2C-D), suggesting an increased trogocytosis. Of note, we found Mef2c, recently shown to restrict neurotoxic phenotype of microglia ^12^, among the downregulated genes; we verified it downregulation in the hippocampus and cortex (Fig S2C-D). Together, these data demonstrated that microglial Cx43 ablation in postnatal development promotes a morphological and transcriptomic change towards a neurotoxic phenotype.

### Developmental cKO of microglial Cx43 promotes neuronal death

Microglia perform multiple roles in developing brain being responsible for clearance of excessive neuronal or glial progenitors by induction of apoptosis or phagoptosis ^3^ ^4^, as well as synapse pruning ^5^ ^36^. We next investigated whether microglial Cx43 ablation-induced neurotoxicity disrupts these developmental processes and examined tissue abnormalities in Cx43^mglcKO^ mice at P7. The most prominent alteration in mice with Cx43 ablation was a profound increase in cell apoptosis marker across different brain regions, as indicated by cleaved Caspase 3 staining (Fig. 3A, C). In addition, we observed numerous cells with pyknosis and karyorrhexis (i.e., condensed, and fractured nucleus), but without cleaved caspase 3 staining, suggesting involvement of alternative cell death pathway(s) (Fig. 3B). Counterstaining with IBA1 and DAPI showed that most cleaved caspase 3 negative dying cells were surrounded by IBA1+ microglial processes (Fig. 3B). These cells may represent targets of microglia-mediated phagoptosis, which is also significantly increased in the cortex and hippocampus following microglial Cx43 ablation (Fig. 3C). We then examined the potential contribution of other cell death pathways. Immunostaining for necrosis marker phospho-MLKL (pMLKL) and pyroptosis marker cleaved GSDMD (cGSDMD) showed no significant changes in P7 Cx43^mglcKO^ mice (Fig. 3G, H). Furthermore, a population of the cleaved Casp3+ cells were positive for Nestin, DCX and NeuN, suggesting the neuronal origin (Fig. 3D-F). Indeed, the number of NeuN+ cells in the hippocampus was reduced by microglial Cx43 ablation (Fig. 3I, Fig. S3A, F). In contrast, the number of Pdgfra+ or Olig2+ oligodendroglial cells was not significantly altered, while the number of GFAP+ astrocytes was increased, possibly due to the increased microgliosis under Cx43 ablation (Fig. 3I, Fig. S3B, C, F). Given that Cx43 cKO microglia show elevated genes involved in complement cascade, implicated in developmental synaptic trimming ^37^, we examined whether synaptic elements were affected in Cx43 cKO mice. However, we did not find significant alteration in the overall level of excitatory synapse marker Vglut1 or inhibitory synapses maker VGAT in different brain regions after ablation of microglial Cx43 (Fig. 3I, Fig. S3D-F).

**Fig. 3.**
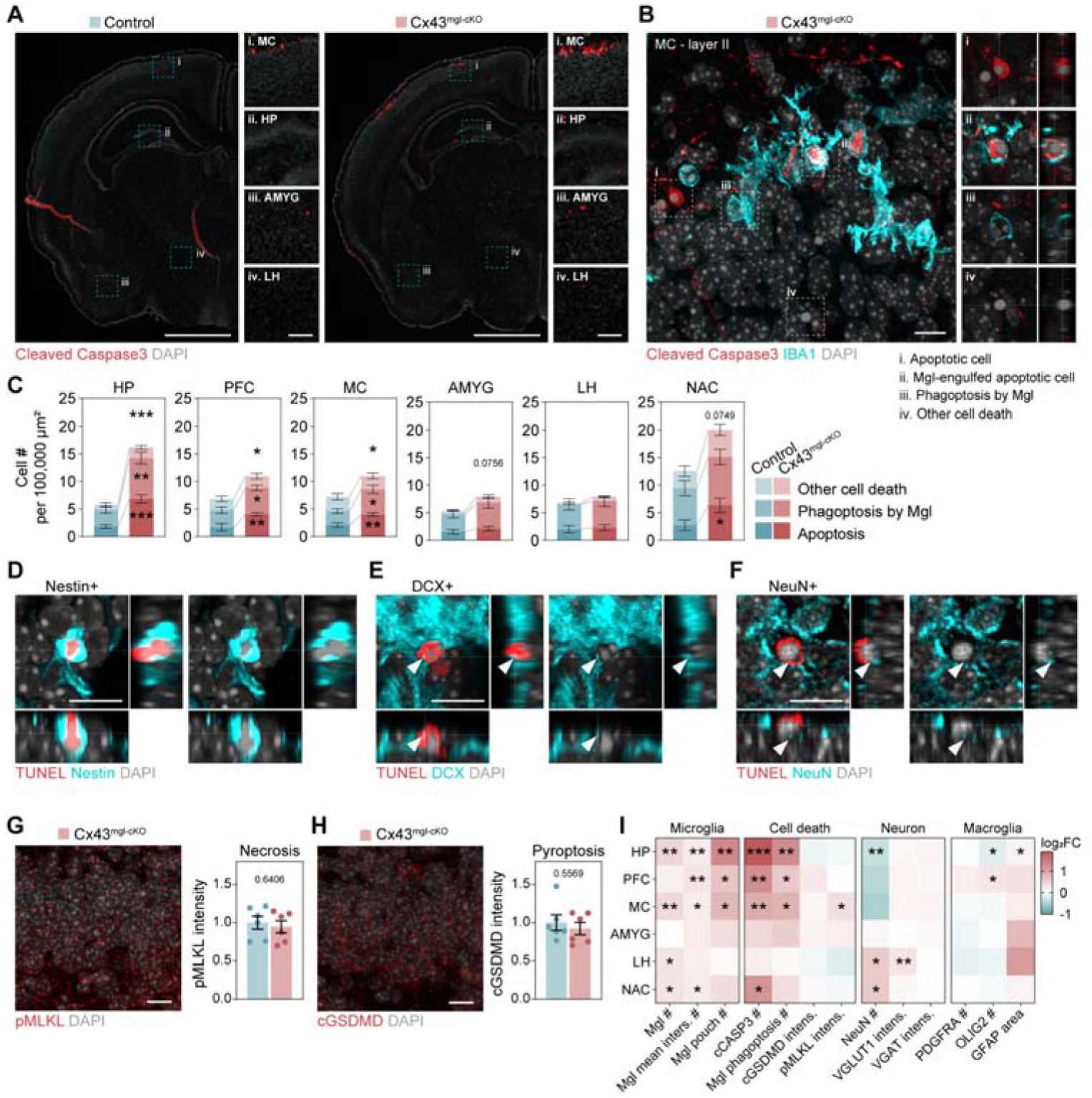
Early postnatal ablation of microglial Cx43 promotes neuronal death. **(A)** Representative images of cleaved-caspase 3 staining in the brains of P7 control and Cx43^mgl-cKO^ mice. High resolution images show microglia in the motor cortex (MC), hippocampus (HP), amygdala (AMYG), and lateral hypothalamus (LH). Scale bar, 500 μm and 50 μm, respectively. **(B)** Representative images of cleaved-caspase 3 and IBA1 fluorescence staining (Scale bar, 10 μm) in Cx43^mgl-cKO^ mouse brains (Scale bar, 10 μm). Orthogonal views of different type of cell death were shown. **(C)** Quantification of the number of apoptotic cells (cCasp3^+^), phagoptosis by microglia, and other cell death, in different brain regions of P7 control and Cx43^mgl-cKO^ mice, N = 6 mice. **(D-F)** Representative images of co-staining for Nestin, DCX and NeuN with Tunel in the Cx43^mgl-cKO^ mouse brain (Scale bar, 10 μm). **(G-H)** Representative images of necrosis marker (phosphor-MLKL, pMLKL) and pyroptosis marker (cleaved-GSDMD, cGSDMD) staining in the brain of Cx43^mgl-cKO^ mice (Scale bar, 10 μm), and quantification of p-MLKL and cGSDMD intensity. N = 6 mice. **(I)** Heatmap of histological abnormalities in P7 Cx43^mgl-cKO^ mice compared to control. See also Fig. S3. N = 6 mice. Plots show mean ± SEM. Significance: \**p* < 0.05, \*\**p* < 0.01, \*\*\**p* < 0.001, \*\*\*\**p* < 0.0001.

We further examined the effect of microglial Cx43 ablation on cell death during embryonic development, given that Cx43 is also highly expressed in embryonic microglia. We induced Cx43 ablation *in utero* by tamoxifen from E14 to E16. At E18, embryonic ablation of microglial Cx43 increased the number of microglia, promoted microglia-mediated phagoptosis, and neuronal apoptosis (Fig. S3G-L). To address the potential background effect of *Cx3cr1^CreERT^* strain ^34^ ^35^, we included the non-Cre littermates (*Cx43^f/+^*or *Cx43^f/f^*) in the analysis, finding that the control group (*Cx3cr1^CreERT/+^*) did not significantly alter the microglia number and neuronal apoptosis compared to the non-Cre group (Fig. S3G, H, J, K). Together, this evidence indicated that developmental ablation of microglial Cx43 promotes neuronal cell death by apoptosis and phagoptosis, suggesting that microglial Cx43 confers a neuroprotective role during embryonic and postnatal development.

### Early postnatal microglial Cx43 ablation caused exhaustion of neurogenic pool and behavioral deficits

We next asked how this hyper-apoptotic phenotype affects later developmental outcomes in Cx43 cKO mice. At P14, the microglia number and morphology were unchanged (Fig. S4A-D), and apoptotic cell number did not differ between control and Cx43^mglcKO^ (Fig. S4E, F). However, Cx43^mglcKO^ mice showed a significant reduction in DCX+ newborn neurons in the dentate gyrus (Fig. S4E, G). Similarly, the number of Ki67+ proliferating neural stem cells was significantly reduced following microglial Cx43 ablation (Fig. S4E, H). We reasoned that in Cx43^mglcKO^ mice, increased apoptosis at the earlier stage (P7) either directly reduced the number of surviving neural stem cells or promoted division of neural stem cells to compensate for the loss of neurons, ultimately leading to premature exhaustion of neural stem cell pool. In Cx43^mglcKO^ mice, reduced neurogenesis in the dentate gyrus persists until P23, as shown by the number of Ki67+ cells and DCX+ immature neurons (Fig. 4A, E, F). The number and morphology of microglia were not altered at this stage either (Fig. 4B-D).

**Fig. 4.**
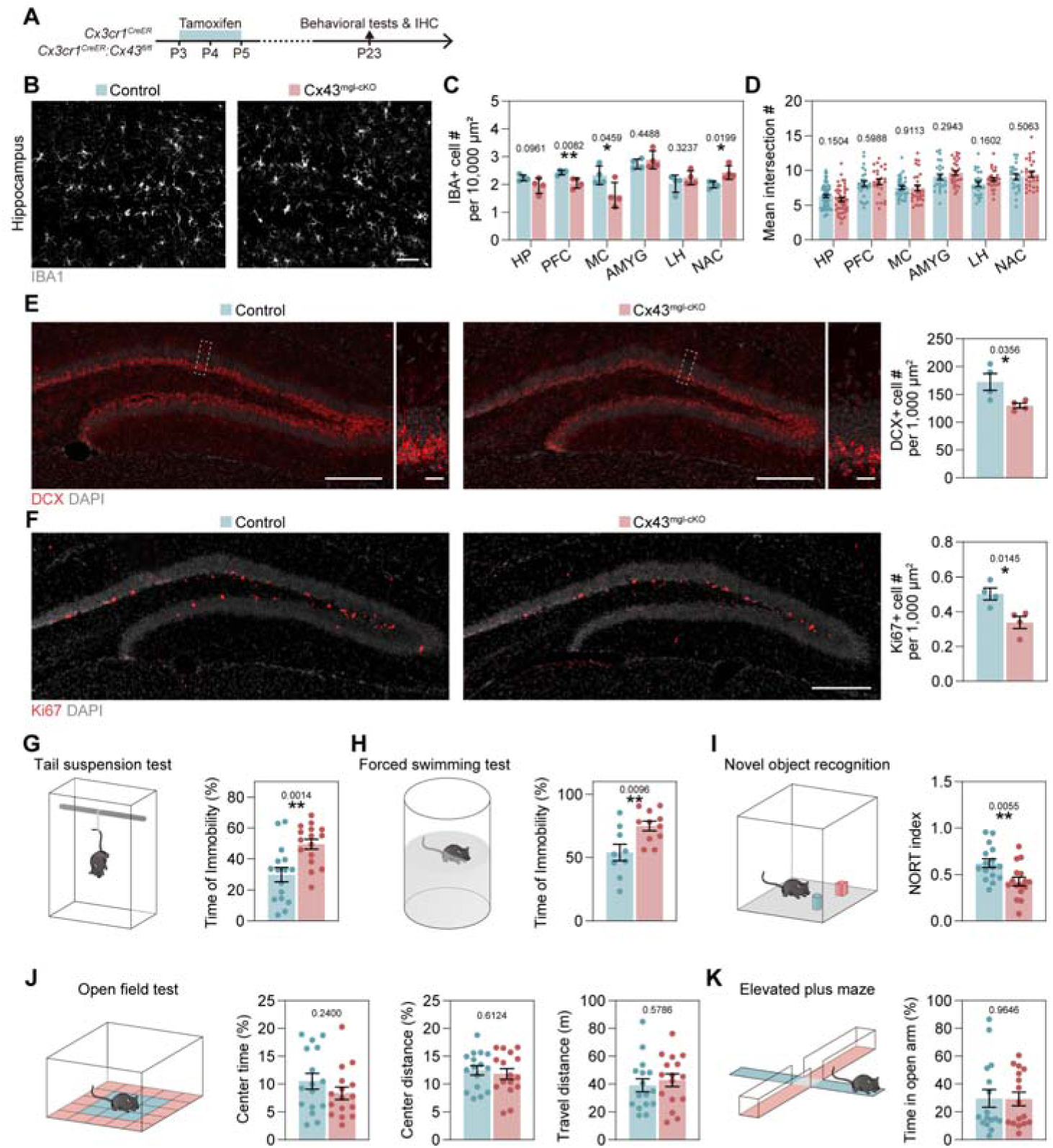
Early postnatal microglial Cx43 ablation caused exhaustion of neurogenic pool and behavioral deficits. **(A)** Schematic of microglial Cx43 knockout induction and examination. **(B)** Representative images of IBA1 staining in the hippocampus from P23 control and Cx43^mgl-cKO^ mice (Scale bar, 50 μm). **(C)** Quantification of IBA1^+^ cells in different brain regions, N = 4 mice. PFC, prefrontal cortex. AMYG, amygdala. LH, lateral hypothalamus. NAC, nucleus accumbens. **(D)** Quantification of mean intersection number from Sholl analysis of IBA1^+^ cell in different brain regions, N = 61, 46 (HP), 27, 24 (PFC), 31, 29 (MC), 25, 27 (AMYG), 23, 27 (LH), 24, 26 (NAC) cells from 4 mice, respectively. **(E)** Representative images of DCX staining in the hippocampus and quantification of DCX+ cells in the dentate gyrus from P23 control and Cx43^mgl-cKO^ mice (Scale bar, 200 μm and 20 μm). N = 4 mice. **(F)** Representative images of proliferation marker (Ki67) staining in the hippocampus and quantification of Ki67+ cells in the dentate gyrus from P23 control and Cx43^mgl-cKO^ mouse hippocampus, staining for (Scale bar, 200 μm). N = 4 mice. **(G)** Immobility time in the tail suspension test. N = 16 mice. **(H)** Immobility time in the forced swimming test. N = 9, 11 mice, respectively. **(I)** NORT index in the novel object recognition test. N = 16 mice. **(J)** Time and distance in the central zone, and total travel distance in the open field test. N = 16 mice. **(K)** Time spent in the open arm in the elevated plus maze test. N = 16 mice. Results are expressed as mean ± SEM. \**p* < 0.05, \*\**p* < 0.01.

Hippocampal neurogenesis give rise to new granule cells that form synapse and integrate into the hippocampal circuit ^38^, and is a critical element of hippocampal synaptic plasticity ^39^. Reduction of postnatal neurogenesis would cause cognitive deficits, depression, and anxiety symptoms ^40^. Thus, we further examined whether postnatal microglial Cx43 ablation may translate into behavioral deficits. Indeed, Cx43^mglcKO^ mice displayed increased immobility time in the tail suspension test and the forced swimming test, suggesting a depressive-like behavior or a passive coping to stress (Fig. 4G-H). During the Novel Object Recognition test, Cx43^mglcKO^ mice showed a decreased preference for the novel object, indicating a cognitive impairment (Fig. 4I). However, Cx43^mglcKO^ mice did not display anxiety trait as determined by the Open Field test and the Elevated Plus Maze test (Fig. 4J-K).

Together, our data suggest that postnatal ablation of microglial Cx43 caused a temporal neurotoxic microglial phenotype, leading to a profound neural cell death. This results in a reduction of neurogenesis in the dentate gyrus, correlating with a depressive-like behavior and cognitive impairment post-weaning.

### Cx43 restricts neurotoxic microgliosis in a neurogenic niche

The temporal neurotoxic microglial phenotype during early postnatal development in Cx43 cKO mice aligned well with their developmental gene expression profile, suggesting a temporal requirement of microglial Cx43 in suppressing neurotoxicity. To examine whether microglial Cx43 ablation in adult mice would cause histological and behavioral defects, we administered tamoxifen to 6-month-old *Cx3cr1^CreERT/+^::Cx43^f/f^* mice for 5 days to induce Cx43 conditional ablation (Fig. S4I). Behavioral tests showed that microglial Cx43 ablation at adulthood caused neither depressive-like behaviors nor cognitive deficits (Fig. S4J-L). This confirmed our previous observation that conditional knockout of microglial Cx43 in 9-month-old wild type mice did not alter the cognitive function ^15,41^. Histological analysis showed that microglial Cx43 ablation in adulthood affected neither the dentate gyrus neurogenesis nor neural cells apoptosis (Fig. S4M-P), which may be due to the low expression of microglial Cx43 in adult brains, suggesting microglial Cx43 is critical for early life brain development.

We further verified whether microglial Cx43 ablation directly contributed to neural cell apoptosis using primary culture. Neural progenitors were isolated from embryonic mice and were either maintained in the neural progenitor state by the addition of FGF and EGF, or allowed to differentiate into neurons. Neural progenitors were co-cultured with primary microglia isolated from control and Cx43^mglcKO^ mice, or treated with control or Cx43 KO microglia-conditioned medium, followed by TUNEL staining to examine cell death (Fig. 5A). Compared to the control group cultured in normal medium, neural progenitors co-cultured with control microglia did not show increased cell death (Fig. 5C). However, co-culturing with Cx43-ablated microglia significantly increased neural progenitor cell death (Fig. 5B, C). Conversely, treatment with conditioned medium from Cx43-ablated microglia did not affect neural progenitor death compared to that from wild-type microglia (Fig. 5C). We deducted that the pro-death phenotype of Cx43-ablated microglia relies on direct interactions with neural progenitors. Indeed, post-mitotic neurons co-cultured with Cx43-ablated microglia did not show increased apoptosis compared to that of wild-type; neither did the treatment with Cx43-ablated microglia-conditioned medium (Fig. 5D). Thus, we hypothesized that Cx43 restricts the neurotoxic phenotype of microglia in the neurogenic niche. To further corroborate this hypothesis, we analyzed microglial alteration in the presence of neural progenitors or neurons. Importantly, Cx43 ablation itself did not alter the number or morphology of microglia in primary culture (Fig. 5E-G). However, the presence of neural progenitors significantly increased the number of microglia with Cx43 ablation, accompanied by an increased cell size, which was not observed in the control microglia (Fig. 5E-G). In contrast, co-culture with neurons showed no effect on microglial number or morphology in either control or Cx43 knockout (Fig. 5E-G). When examining microglial morphology, we noticed that microglia co-cultured with neural progenitors extended IBA1-high lamellipodia (a structure that resembled the phagocytic pouch observed *in vivo)* that contacted viable or TUNEL+ neural progenitors (Fig. 5H-I). Together, Cx43 ablation unleashes a contact-dependent neurotoxic microglial phenotype that triggers apoptosis of neural progenitors but not neurons, and is accompanied by microglial accumulation, cellular hypertrophy, and lamellipodial contacts with dying progenitors.

**Fig. 5.**
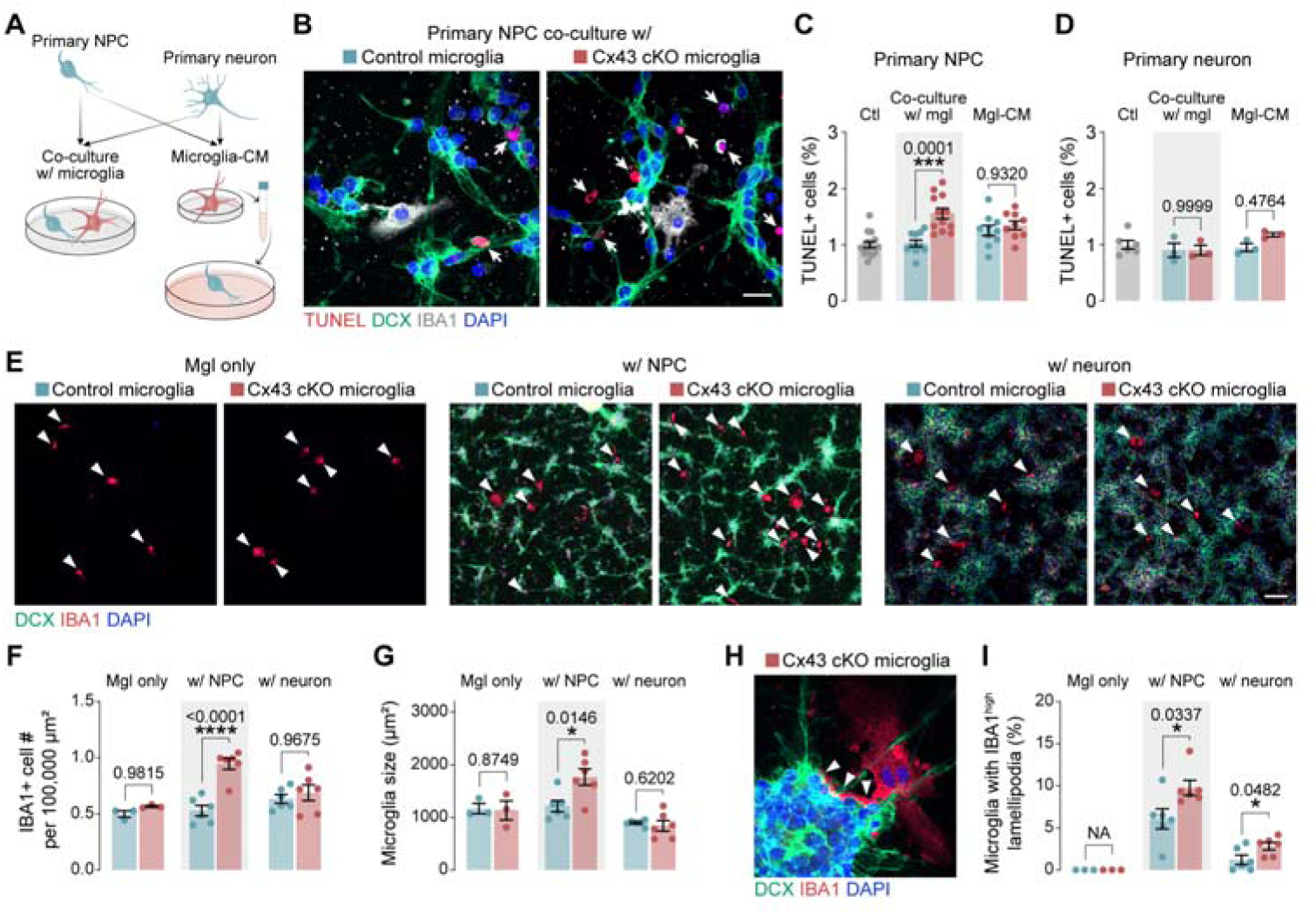
Cx43 restricts neurotoxic microgliosis in a neurogenic niche. **(A)** Schematic of primary cell culture design. **(B)** Representative images of primary neural progenitor cell (NPC) co-cultured with control and Cx43 cKO microglia, staining for DCX and IBA1 after in situ TUNEL labeling (Scale bar, 50 μm). Arrows highlight TUNEL+ cells. **(C)** Quantification of TUNEL^+^ cell percentage in primary NPC cocultured with microglia, or with microglia conditioned medium (CM). N = 14 (control), 10, 12 (microglia-co-culture), 10, 9 (microglia-CM) coverslips. **(D)** Quantification of TUNEL^+^ cell percentage in primary neuron cocultured with microglia, or with microglia conditioned medium (CM). N = 6 (control), and 3 (microglia-co-culture or microglia-CM) coverslips. **(E)** Representative fluorescence microscopy images of primary control and Cx43 cKO microglia with or without NPC or neuron co-culture, staining for DCX and IBA1 (Scale bar, 100 μm). Arrowheads highlight IBA1+ microglia. **(F-G)** Quantification of microglia number and cellular area when cultured alone or with NPC or neuron co-culture. N = 3 (microglia only) or 6 (co-culture) coverslips. **(H)** Representative images of primary Cx43 cKO microglia with NPC co-culture (Scale bar, 50 μm). Arrowheads highlight the IBA1^high^ lamellipodia contacting NPC. **(I)** Quantification of microglia with IBA1^high^ lamellipodia when cultured alone or with NPC or neuron co-culture. N = 3 (microglia only) or 6 (co-culture) coverslips. Results are expressed as mean ± SEM. Significance: \**p* < 0.05, \*\*\**p* < 0.001, \*\*\*\**p* < 0.0001.

### Microglial Cx43 hemichannel did not confer neuroprotective role in development

Microglial Cx43 hemichannels are known to be operational mainly in pathology, being implicated in the release of factors interfering with neuronal viability ^15^. However, this neurotoxic functional mechanism of microglial Cx43 is in direct contrast to our observation in development. We investigated Cx43 hemichannel activity in developing brains by using dye uptake protocol on acute brain slices from mice at P1, P7, P14, and P21. Microglia at P1 and P7 display higher levels of dye uptake compared to those at P21; dye uptake was suppressed by specific Cx43 hemichannel inhibitor TAT-Cx43_266-283_ (Fig. S5A), indicating that microglial Cx43 hemichannels are active during development.

We therefore examined whether inhibition of microglial Cx43 hemichannels mimics the neurotoxic phenotype observed in the Cx43 conditional knockout. To this end, we administrated a Cx43 hemichannel specific inhibitor, TAT-Cx43_266-283_ peptide (TAT-Cx43) ^15,41^, to wild type mouse pups once a day from P2 to P7, with TAT peptide as the control (Fig. S5B). TAT-Cx43_266-283_ peptide was able to cross the blood brain barrier and interact with microglial Cx43 ^15^. Blockade of hemichannels did not phenocopy the histological changes triggered by in microglial Cx43 ablation. Histological analysis on P7 revealed that TAT-Cx43_266-283_ treatment significantly reduced the number of microglia in the hippocampus, in contrast to that of Cx43^mgl-Cko^ mice, but did not affect the microglia morphology (Fig. S5C, D). The expression of FCGR3 and C1q was also reduced by TAT-Cx43_266-283_ treatment, yet the expression of CLEC7A or MEF2C was not altered (Fig. S5E, F). We further examined whether apoptosis was affected by TAT-Cx43_266-283_ treatment and found that the number of cleaved Caspase3+ cells in the hippocampus was significantly reduced (Fig. S5G).

The number of TUNEL+ cells, however, remained unchanged (Fig. S5H). The TAT-Cx43_266-283_ did not affect the numbers of DCX+ and Ki67+ cells in the dentate gyrus when examining on P14 (Fig. S5I-K). These data suggested that Cx43 hemichannels in developing microglia promotes a proliferative, phagocytic phenotype; however, inhibition of Cx43 hemichannel by TAT-Cx43_266-283_ treatment could not phenocopy the outcome of microglial Cx43 conditional knockout. Therefore, a non-canonical function of microglial Cx43 might be at play.

### Nucleus-localized Cx43 limits neurotoxic microglia in perinatal development

Considering the nuclear-localization of Cx43 in developmental microglia, we investigated whether nuclear Cx43 participates in the regulation of neurotoxic phenotype. We used an AAV11-mIBA1-miR9T expression vector targeting microglia to overexpress Cx43-HA and a SV40-NLS conjugated Cx43-HA in hippocampal microglia. We first confirmed the microglial specificity of this viral tool, as off-target transduction has been reported in several brain regions. Here, however, the virus specifically targeted microglia in the developing hippocampus when injected at P2 (99.34 ± 0.23%), with 55.96 ± 1.53% of hippocampal microglia successfully transfected (Fig. S6A-B). We then used this system to overexpress wild-type mouse Cx43 and NLS-conjugated Cx43 in microglia. Immunostaining of the HA tag confirmed the successful expression of Cx43 in microglia. Compared to the wild-type Cx43, the NLS-Cx43 displays a significantly higher percentage of nucleus-localization (Fig. 6A-C). The overexpression of wild-type Cx43 did not affect microglial morphology; however, the overexpression of NLS-Cx43 significantly reduced the ramification of microglia (Fig. 6D). In addition, overexpression of NLS-Cx43, but not wild type Cx43, significantly reduced the expression of CLEC7A and FCGR3 (Fig. 6E, F). Concomitantly, the number of microglia phagocytic pouch was also reduced in NLS-Cx43 overexpression (Fig. 6G). We further examined whether overexpression of Cx43 in microglia affects neural apoptosis. Labeling with cleaved caspase-3 showed that while wild type Cx43 did not alter the number of apoptotic cells, the NLS-Cx43 overexpression significantly reduced apoptosis in the hippocampus (Fig. 6H, I). Together, overexpression of nuclear-localized Cx43 in perinatal hippocampal microglia reduced their ramification, lowered neurotoxic marker expression and phagocytic pouch formation, and decreased neuronal apoptosis, suggesting a critical role of nucleus-localized Cx43 in restricting microglial neurotoxic phenotype in development.

**Fig. 6.**
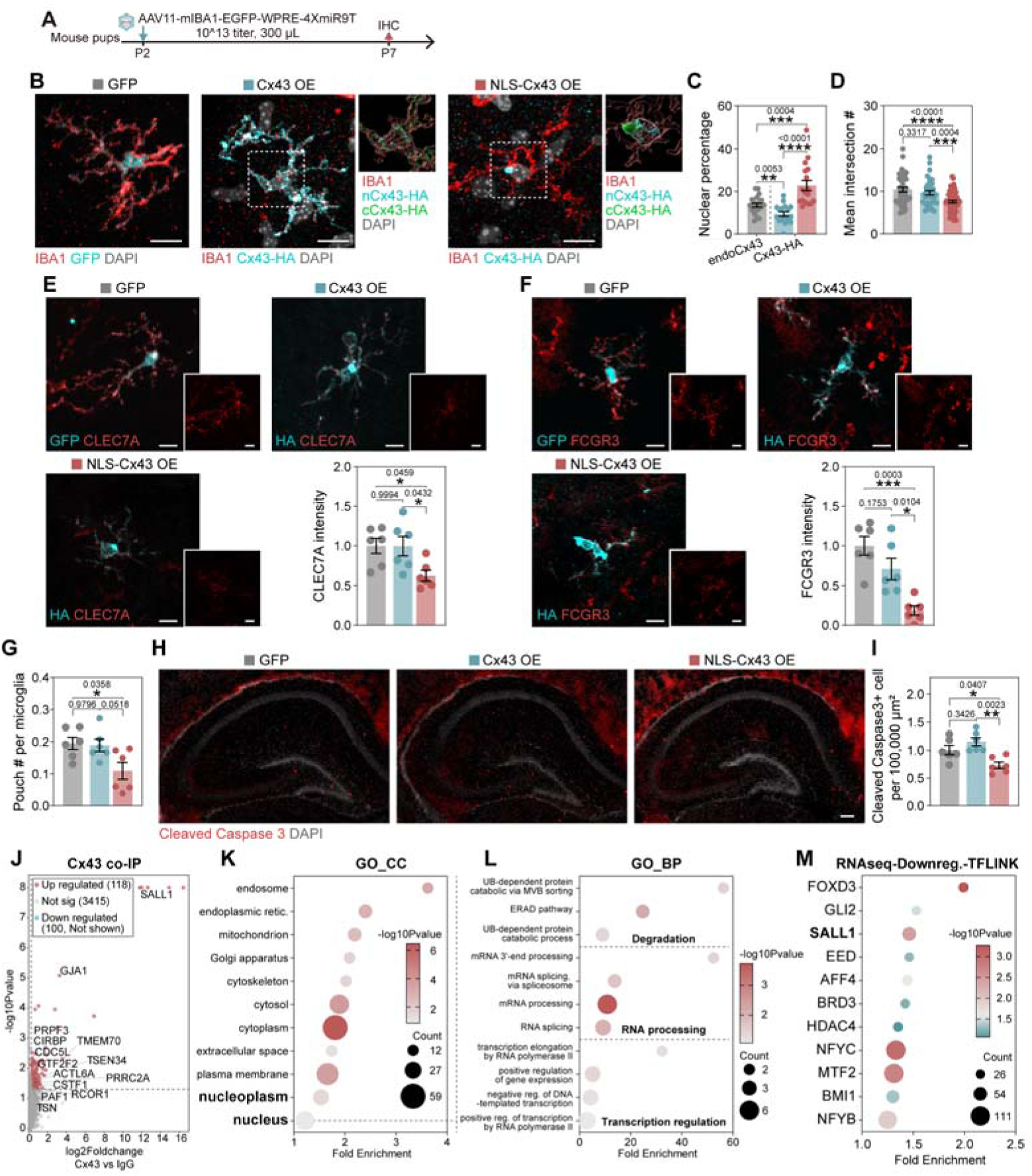
Nucleus localized Cx43 limits neurotoxic microglia in perinatal development. **(A)** Schematic of AAV-induced Cx43 overexpression experiments. **(B)** Representative images of AAV transfected microglia in hippocampus, staining for HA, GFP, and IBA1, with 3D reconstruction of IBA1/DAPI masked Cx43-HA (Scale bar, 10 μm). nCx43-HA, nucleus Cx43-HA. cCx43-HA, cytoplasmic Cx43-HA. **(C)** Quantification of nuclear percentage of endoCx43 and Cx43-HA, NLS-Cx43-HA. endoCx43, endogenous Cx43 in GFP-transfected microglia. N = 21, 17, 17 cells from 3 mice. **(D)** Quantification of mean intersection number from Sholl analysis of transfected microglia using IBA1 staining. N = 21, 17, 17 cells from 3 mice. **(E)** Representative images of transfected microglia staining for CLEC7A and GFP/HA (Scale bar, 10 μm). CLEC7A intensity of transfected microglia was quantified. N = 6 mice. **(F)** Representative images of transfected microglia staining for FCGR3 and GFP/HA (Scale bar, 10 μm). FCGR3 intensity of transfected microglia was quantified. N = 6 mice. **(G)** Quantification of pouch number per transfected microglia. N = 6 mice. **(H)** Representative images of transfected hippocampus staining for cleaved Caspase 3 (Scale bar, 50 μm). **(I)** Quantification of cleaved Caspase 3^+^ cells in hippocampus. N = 6 mice. **(J)** Volcano plot of Cx43 co-IP experiment, with normal IgG as control. Cx43 (GJA1) and nucleus-localized proteins were highlighted. **(K)** Functional annotation (Gene ontology_cellular compartment) of Cx43 co-IP enriched proteins. **(L)** Functional annotation (Gene ontology_biological process) of nucleus proteins co-IPed with Cx43. **(M)** TFLINK enrichment of genes downregulated in Cx43 cKO microglia revealed by RNA-seq. Results are expressed as mean ± SEM. Significance: \**p* < 0.05, \*\**p* < 0.01, \*\*\**p* < 0.001, \*\*\*\**p* < 0.0001.

We next asked what the functional mechanism of nucleus-localized Cx43 is. One of the hypothetic mechanisms is Cx43 interaction with chromatin to regulate transcription. To this end, we analyzed interaction between Cx43 and chromatin using ChIP-seq. We verified the immunoprecipitation capacity of Cx43 antibody (Fig. S6C). However, no significant enrichment was observed in the ChIP-seq (Fig. S6D). Cut-Tag and cut-run assays did not yield confident, repeatable peak signals either (Fig. S6D). These results suggest that Cx43 may not interact with chromatin, or the Cx43 antibody we used may not be suitable for these assays. Nevertheless, we examined the protein interactome of Cx43, using Cx43 co-IP and mass spectrometry analysis on acute-isolated P7 wild type mouse microglia. We found GJA1 (Cx43) among the co-IPed proteins, together with several reported Cx43 binding partners, such as YWHAE (14-3-3ε) ^42^, EZR ^43^, and TSG101 ^44^, confirming the successful Cx43 pull-down (Fig. 6J, Fig. S6E-G). Gene ontology cellular compartment (GO_CC) annotation showed that numerous nucleus and nucleoplasm proteins co-immunoprecipitated with Cx43, suggesting possible interactions (Fig. 6K, L). Gene ontology-biological process annotation showed that these nuclear proteins were involved in transcriptional regulation (e.g., SALL1, PAF1, GTF2F2, etc.), RNA processing (e.g., PRPF3, PTBP3, CDC5L, etc.), as well as ubiquitin-dependent protein degradation (e.g., UBQLN1, UBE4A, UBXN4, etc.) (Fig. 6L). When using TFLINK to predict the transcription factors of down-regulated genes in Cx43 knockout microglia, we found SALL1 among the plausible transcription factors at play (Fig. 6M). SALL1 is a microglial enriched transcription factor, which has been shown to suppress proinflammatory microgliosis to maintain hippocampal neurogenesis ^45^ ^46^. Thus, nucleus Cx43 may act synergistically with SALL1 to restrict neurotoxic microgliosis during development.

### Dynamic nucleus translocation of Cx43 in developmental microglia

We revealed the critical role of nucleus Cx43 in limiting neurotoxic microglial phenotypic change during development. We then asked whether microglial Cx43 nucleus translocation is dynamically regulated during development. We first sought to examine the nuclear localization signal (NLS) and nuclear export signal (NES) in Cx43. cNLS mapper ^47^ and LocNes tool ^48^ predicted several importin-α-dependent NLSs and classical NESs in Cx43, respectively (Fig. 7A). Among the NLSs, two regions raised our attention (aa237-290, NLS1; and aa343-373, NLS2) due to: (i) they are located at the flexible C-terminus of Cx43, and (ii) numerous post-translational modification sites are found in these two regions ^49^, which may participate in the dynamic nucleus transportation of Cx43. We overexpressed GFP fused to the N-terminus of these two putative Cx43 NLS in primary microglia (Fig. 7B, Fig. S7A-B). Compared to GFP control, only NLS2-GFP (but not NLS1-GFP) show significantly higher nucleus localization (Fig. 7C, Fig. S7A, C). As a complementary approach, we overexpressed full length Cx43 and its NLS truncated mutants (Fig. B). Deletion of NLS2 significantly reduced the percentage of Cx43 translocated into the nucleus (Fig. 7D, Fig. S7A, D). Simultaneous deletion of NLS1 and NLS2 further impeded the nucleus translocation of Cx43 (Fig. 7D, Fig. S7A, D), suggesting a co-operative role of these two NLSs.

**Fig. 7.**
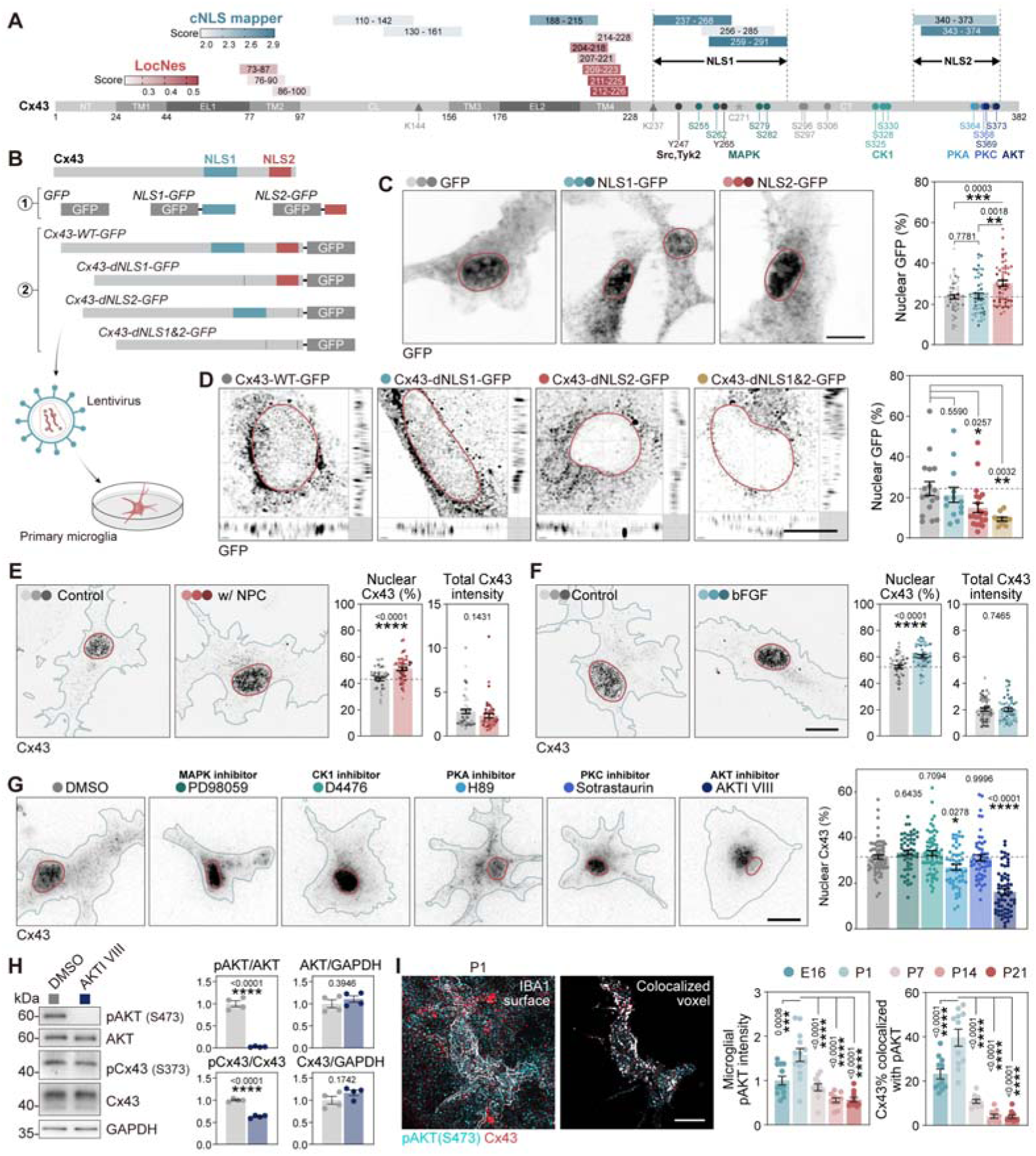
Dynamic nucleus translocation of Cx43 in developmental microglia. **(A)** Schematic of Cx43 protein domains, cNLS mapper predicted NLS, locNes predicted NES, and reported post-translational modifications and binding partners ^49,76^. Circle, phosphorylation. Triangle, sumoylation. Asterisk, nitrosylation. **(B)** Construct of Cx43 NLS1- and NLS2-GFP fusion proteins, and Cx43-dNLS mutant-GFP fusion proteins, which were packaged in lentivirus and transfected into primary microglia. **(C)** Representative images of NLS-GFP overexpressed microglia (Scale bar, 10 μm). Percentage of nucleus-localized GFP was quantified. N = 50 cells from 3 experiments. **(D)** Representative orthogonal images of wildtype Cx43 and NLS-deleted Cx43 mutants overexpressed microglia (Scale bar, 10 μm). Percentage of nucleus-localized GFP was quantified. N = 17, 13, 21, 10 cells from 3 experiments. **(E)** Representative images of primary microglia with or without neural progenitor cell Transwell co-culture, staining for Cx43 (Scale bar, 10 μm). Nuclear Cx43 % and total Cx43 intensity were quantified. Plot shows mean ± SEM. N = 52 cells from 3 experiments. **(F)** Representative images of primary microglia with or without bFGF treatment (Scale bar, 10 μm). Nuclear Cx43 % and total Cx43 intensity were quantified. Plot shows mean ± SEM. N = 52, 49 cells from 3 experiments. **(G)** Representative images of primary microglia treated with MAPK, CK1, PKA, PKC, AKT inhibitors, staining for Cx43. (Scale bar, 10 μm). Nuclear Cx43 % was quantified. N = 65, 53, 65, 51, 54, 63 cells from 3 experiments. **(H)** Western blot analysis of primary microglia lysates, blotting with pAKT (S473), AKT, pCx43 (S373), Cx43, GAPDH. Plot shows mean ± SEM. N = 4 experiments. **(I)** Representative images of P1 mouse hippocampal microglia staining for IBA1, pAKT (S473) and Cx43, with IBA1 reconstructed surface shown (Scale bar, 10 μm). Right image shows colocalized voxels between pAKT (S473) and Cx43. The microglial pAKT intensity and the percentage of Cx43 signal with pAKT colocalization were quantified in E16, P1, P7, P14 and P21 mouse microglia. N = 12, 13, 11, 10, 10 cells from 3 mice for pAKT intensity, N = 11, 11, 10, 10, 10 images from 3 mice for colocalization test. Results are expressed as mean ± SEM. Significance: \**p* < 0.05, \*\**p* < 0.01, \*\*\**p* < 0.001, \*\*\*\**p* < 0.0001.

Microglial Cx43 nucleus translocation was prominent at perinatal stages, when abundant number of neural progenitors and immature neurons was present. Thus, we wondered if neural progenitor-derived factors could promote Cx43 nucleus translocation. To address this, we cocultured primary microglia with or without neurospheres in a Transwell setting. The presence of neurospheres promotes the translocation of Cx43 to the nucleus in microglia (Fig. 7E, Fig. S7E). Neural progenitor release factors regulating microglia phenotypes, including bFGF ^7^. We found that addition of bFGF to the culture medium also promoted Cx43 nucleus translocation (Fig. 7F, Fig. S7F). In contrast, pathological or inflammatory agents, including TNFα, LPS, and Aβ, reduced the percentage of nucleus localized Cx43 (Fig. S7F). These results suggest that microglial Cx43 nucleus translocation is dynamically regulated, and is promoted in the neurogenic niche during development.

As we demonstrated above Cx43 nuclear translocation could be promoted by co-culturing with neural progenitor cells or exposure to bFGF. Both treatments could activate various kinases that might be involved in the Cx43 translocation. Notably, Cx43 was reported to have numerous phosphorylation sites of these kinases, including MAPK, CK1, PKA, PKC, and AKT, which impact on Cx43 localization and function ^49^. We sought to dissect which of these kinases might be involved in the translocation of Cx43 by treatment with appropriate inhibitors in primary culture. We found that inhibition of AKT by AKTI-III substantially reduced nuclear Cx43 (Fig. 7G, Fig. S7G). Inhibition of PKA with H89 also reduced nuclear Cx43, but to a lesser extend (Fig. 7G, Fig. S7G). Both AKT and PKA can phosphorylate Cx43 at S369/S373 and S364, respectively, all located within the putative NLS2 of Cx43 (Fig. 7A), further consolidating NLS2 role in directing Cx43 nuclear translocation. Treatment with AKTI-III efficiently eliminated AKT phosphorylation (S473), and reduced Cx43 S373 phosphorylation by 37.06 ± 2.29% (Fig. 7H). Immunostaining of IBA1 and phospho-AKT (S473) in developing brain sections showed that activation of microglial AKT pathway is more prominent in perinatal period (P1) (Fig. 7I). Also, significantly higher percentage of microglial Cx43 colocalized with phosphor-AKT (S473) at this stage (Fig. 7I).

Together, these results provide a mechanistic blueprint of how Cx43 nucleus translocation is regulated in developing microglia. Neurogenic niche □ derived signals such as bFGF promote AKT □ dependent phosphorylation of Cx43 at S373 within its C □ terminal NLS, facilitating subsequent nucleus translocation.

## DISCUSSION

Here, we identify a previously unrecognized nuclear function of full-length Cx43 in developmental microglia, revealing that this canonical membrane protein also acts in the nucleus to restrain neurotoxic microglial activation during brain development. We show that nuclear accumulation of multimeric Cx43 is a transient developmental feature in both human and mouse microglia, is promoted by neurogenic niche-derived signals such as bFGF, and depends on AKT-driven phosphorylation. Functionally, loss of microglial Cx43 during early postnatal life, but not adulthood, drives a neurotoxic programme characterized by excessive phagoptosis and neuronal apoptosis, depletion of the hippocampal neurogenic pool, and lasting depressive-like and cognitive deficits, whereas hemichannel blockade fails to phenocopy these effects. Together, these findings provide an explanation for a previously unknown non-canonical function of Cx43 that is critical for maintaining a protective microglial state and safeguarding normal brain development.

Microglia enter the brain parenchyma during early embryonic development, and reciprocally interact with the brain cellular elements, which is critical for the proper development of CNS ^1^ ^10^. During this process, microglial phenotype is dynamically remodeled by the neurogenic niche through neurotropic factors such as VEGF, which promote microglia proliferation and phagocytic activity ^7^. In the fetal cortex, microglia actively engulf neurons ^3^ and OPCs ^4^, remodel synapses ^5^ and myelin ^6^, and maintain the structural integrity at the fetal cortical boundaries by phagocytosis of extracellular matrix ^50^. Disruption of these processes results in an aberrant developmental programming and the subsequent neurological and behavioral disorders ^6^ ^8^ ^9^. Microglial phagocytosis of dying cells and cell debris can trigger neurotoxic microgliosis as seen during neuronal injury and neurodegeneration ^10^ ^11^. However, in development microglia seem to possess a program to adjust such phenotypic change. For instance, MEF2C halts the neurotoxic microgliosis during development ^12^. Here we found that microglial Cx43, in response to neurotrophic factors, serves as a brake on the neurotoxic phenotype in the developing CNS: niche-derived signals such as bFGF promote its nuclear translocation, where Cx43 interacts with transcriptional regulators like SALL1 to restrain proinflammatory gene expression. This nuclear checkpoint thus tempers the microglial response to developmental cell death, preventing excessive phagoptosis and preserving the neurogenic pool. Loss of this brake, as demonstrated by our conditional knockout, unleashes a self-amplifying neurotoxic state that echoes degenerative processes, yet is uniquely developmentally restricted.

Nuclear localization of Cx43 has rarely been reported *in vivo*. A truncated form of Cx43 was found to in the nucleoplasm, regulating transcriptional activation ^23^ ^24^. Full-length Cx43 has also been localized in the nuclear envelope ^25^ ^26^, forming functional channels ^25^, or engaging with DNA damage response complexes ^26^. Several signaling pathways can regulate Cx43 nuclear import, including Wnt and PKA ^25^. However, the physiological relevance was unclear before this study. Here, we describe a different form of Cx43 nucleus localization: the full-length Cx43 in the nucleoplasm. Notably, the nuclear accumulation of full-length Cx43 displays a cell-type and developmental specificity. It is selectively enriched in microglia compared with other brain parenchymal cells, and within the microglial population, it is substantially higher during the perinatal period than in juvenile or adult stages. We discover a novel key mechanism mediating the dynamic microglial Cx43 nuclear import during neurodevelopment. In the neurogenic niche, neurotrophic factor bFGF activates AKT signaling, which in turn phosphorylates Cx43 at C-terminus. AKT phosphorylation of Cx43 could promote docking of 14-3-3 onto Cx43 c-terminus ^42^, which may facilitate protein nucleus translocation ^51^. Other, yet unknown pathways may also promote Cx43 nucleus translocation, which requires further systemic analysis.

While we showcased several cellular cascades and signaling pathways contributing to full-length Cx43 nucleus translocation, it remains difficult to fathom how a transmembrane protein such as Cx43 could be extracted from the cellular membrane and enter the nucleoplasm. In general, nuclear translocation of transmembrane protein is thought to involve proteolytic cleavage of the cytoplasmic domain by enzymes such as γ-secretase, thereby releasing an intracellular fragment that can be imported into the nucleus, as exemplified by Notch receptor signaling ^52^. However, in addition to Cx43, several transmembrane proteins have also been reported to translocate into the nucleoplasm in their full-length form and interact with chromatin; these include including epidermal growth factor receptor (EGFR) ^53^, nerve growth factor receptor (NGFR) ^54^, platelet-derived growth factor receptor (PDGFR) ^54^, fibroblast growth factor receptors (FGFRs) ^55^, growth hormone receptor ^56^, insulin-like growth factor receptors ^57^, Transferrin receptor 1 ^58^, and CD44 ^59^. Despite these observations, the underlying mechanism remains poorly defined. It has been proposed that this process may occur in conjunction with nuclear import pathways ^58^. In the case of EGFR, the translocon Sec61β has been suggested to facilitate its trafficking across the inner nuclear membrane and subsequent release into the nucleoplasm ^60^. Alternatively, membrane proteins may first undergo endosomal escape into the cytosol before nuclear transportation ^57^. For Cx43 specifically, interaction with the heat shock protein^61^, or involvement in phase separation process^62^, might facilitate its escape from membrane. Further studies are required to delineate the precise mechanisms governing these processes.

Aberrant nuclear translocation of Cx43 may be linked to pathophysiology of neurodevelopmental and neuropsychiatric diseases. Mutations in *GJA1* give rise to oculodentodigital dysplasia (ODDD), a pleiotropic disease which can manifest as neuropsychiatric disorders characterized by mood instability and cognitive impairment ^63^. These mutations of Cx43 could cause disease relevant aberrant subcellular distribution. For instance, A40V mutant was found in the nucleus ^64^. Some other Cx43 mutants including G21R, G138R, and fs230 also seems to show nucleus translocation ^65^ ^66^. While the reduction of gap junction channel function of these mutants might be responsible for ODDD-related neuropsychiatric disorders ^64^ ^65^, dysregulated Cx43 translocation and non-canonical Cx43 functions could be also involved. Cx43 is expressed not only in developing microglia but also in neural progenitors, astrocytes, and ependymoglia. Therefore, aberrant nuclear translocation of Cx43 and its non □ canonical functions could potentially affect multiple cell types, not just microglia, and thereby contribute to the neuropsychiatric features of ODDD.

Beyond its canonical function as a plasmalemmal channel protein, Cx43 exhibits diverse non-canonical roles linked to its alternative subcellular localization. Notably, Cx43 localizes to the inner mitochondrial membrane in a HSP90-dependent manner, where it regulates mitochondrial respiration, ATP production, potassium influx, and reactive oxygen species generation ^20^ ^21^. Cx43 also contributes to intercellular adhesion independent of its channel function, serving as a mechanical coupling molecule essential for cell morphology ^17^. Additionally, the C-terminus of Cx43 acts as a scaffolding platform, regulating the activity of several kinases, including Akt, Erk, and Yap ^18^ ^19^. Cx43 further facilitates long-distance intercellular communication by localizing to extracellular vesicles ^22^, enabling the transfer of cargo including mitochondria between cells ^67^. Truncated isoforms, such as GJA1-20k, participate in trafficking full-length Cx43 to the plasmalemmal ^68^, maintain mitochondrial homeostasis ^69^, and regulate transcription ^24^. Despite these advances, our understanding of the non-canonical function of Cx43 remains largely compartment-centric, with most studies confined to a single organelle. Consequently, the distribution of Cx43 across subcellular compartments, the mechanisms governing its dynamic redistribution, and the physiological consequences of such compartmentalization remain poorly defined. In developing perinatal microglia, our finding suggests that plasmalemmal Cx43 and nuclear Cx43 serve distinct functions, with plasmalemmal Cx43 hemichannel activity showing neurotoxic potential and nuclear Cx43 exerting a prominent neuroprotective role. However, the contribution of Cx43 in mitochondria and other subcellular compartments remains unresolved, as does the extent to which loos of each pool contributes to the neurotoxic phenotype in the conditional knockout. Clarifying these compartment-specific roles will be important for determining whether therapeutic strategies should broadly modulate Cx43 or selectively target specific subcellular pools while preserving protective functions.

In summary, we identify a nuclear function of microglial Cx43 that restrains neurotoxic gene programs and protects the developing brain, extending connexin biology beyond channel activity to include a transcriptional co-regulation. Key questions remain, including whether Cx43 directly engages chromatin, the composition of its nuclear interactome, and whether similar translocation occurs in other cell types. Defining how Cx43 is positioned across subcellular compartments in health and disease may inform compartment-specific therapeutic strategies that preserve its neuroprotective nuclear role while limiting pathogenic hemichannel activity in neurodevelopmental and neuropsychiatric disorders.

## METHODS AND MATERIALS

### Human samples

Human embryonic brain tissues were obtained from the School of Medicine, Sun Yat-sen University, under the ethical approval of the School of Medicine, Sun Yat-sen University Institutional Review Board (NO. 2026-27). Informed consent was obtained by participants. Tissues were fixed in formalin for 2 weeks, and dehydrated in 30% Sucrose in PBS for 2 days before embedding for cryosections at 20 μm. For immunostaining, the sections were processed for citric acid antigen retrieval, permeabilized with 0.5% Triton-X100 in PBS (0.5% PBST) for 1 hour, blocked with 2% BSA in PBST for 1 hour, and incubated with primary antibody overnight at 4 □, followed by secondary antibody, counterstained with DAPI, and mounted for microscopic observation. Primary antibodies and dilution are listed below: Rabbit-anti-Cx43 (Merck, C6219, 1:200), Goat-anti-IBA1 (Abcam, ab5076, 1:400).

### Animals

Mice were maintained in a PC2 pathogen-free animal facility in a 12 h/12 h light/dark cycle with a temperature of ∼20 °C and ∼50% humidity. All procedures were performed under the ethical approval of the Sun Yat-Sen University Institutional Animal Care and Use Committee (NO. SYSU-IACUC-2024-000452). Cx3cr1^CreERT^ mice, as previously reported, were obtained from the Jackson Lab (Strain # 21160). They were mated with Cx43^flox^ (Strain # 008039) mice to generate *Cx3cr1^CreERT^::Cx43^flox^* mice (APP/PS1:Cx43^mgl-cKO^). The *Cx3cr1^CreERT/+^::Cx43^+/+^* and *Cx3cr1^CreERT/+^::Cx43^f/+^*littermates were used as controls. To induce Cx43 knockout, 10 mg/mL tamoxifen in ethanol/corn oil mixture (1:9, 50 μL) was given to mice via gavage once daily for 3 consecutive days starting from P3, or from E14 for embryonic knockout. In adult mice, 30 mg/mL tamoxifen was given to mice via gavage at 50 μL each day for 5 consecutive days. For TAT-Cx43_266-283_ treatment, mice received TAT or TAT-Cx43_266-283_ (4 ng/g) injection once per day from P2 to P7. Mice of both sexes were used.

### Behavioral tests

Before behavioral tests, mice were gently handled for 3 consecutive days by the experimenter. Tests were conducted in the following sequence: Open Field Test, Novel Object Recognition, Elevated Plus Maze, Tail Suspension, and Forced Swimming Test. Videotaping and data analysis were conducted using the VisuTrack software and hardware (Xinrun, Shanghai).

**Open Field Test** was used to determine the anxiety-like behaviors and locomotor function. Mice were placed in the center of the arena (50 × 50 × 40 (height) cm^3^) and were allowed to explore for 20 min.

**Novel Object Recognition test** was used to determine the cognitive function. In the familiarization session, mice were placed in a 25 × 25 × 40 (height) cm^3^ white box with 2 rectangular plastic objects placed 8 cm away from the walls and allowed to explore for 5 min. The test session took place two hours later, where one of the rectangular objects was replaced by a cylinder (the novel object) before the mouse was re-introduced into the apparatus. The discrimination index calculated as (Time^new^-Time^old^) / (Time^new^-Time^old^) was used to determine the exploratory preference toward the novel object.

**Elevated Plus Maze** was used to determine the anxiety trait. Mice was placed in the elevated plus maze with two opposing closed arms and two opposing open arms. Mouse activity was recorded for 10 min. The time spent in the open arm were measured.

**Tail Suspension test** was used to determine the depressive trait. Briefly, mice were suspended by sticking the end of the tail to the edge of a shelf using adhesive tape for 6 min. Mouse activity was recorded and the immobility time was measured.

**Forced Swimming test** was used to determine the depressive trait. Briefly, mice were placed in an inescapable cylindrical transparent tank (30 cm height × 20 cm diameter) that was filled with water and their escape-related mobility behavior was recorded for 6 min. The immobility time of the last four minutes was analyzed.

### Microglia isolation

For primary culture and RNA-seq, microglia were isolated using immunopanning as previously described with minor modifications. Briefly, postnatal day 7 mouse pups were subjected to hypothermia-induced anesthesia, and euthanized by rapid decapitation with sharpened scissors. The brains were isolated, minced, and digested with 1 mg/mL papain for 1 hour at 37 □ . Tissues were then triturated with 10 mL pipette, and the resulting cell suspension was passed through a 70 μm cell strainer, and subjected to immunopanning with the rat-anti-mouse CD45 antibody (BD, 553076, 1:200) and the secondary goat-anti-rat antibody (Jackson ImmunoResearch, 112-005-003) to isolate microglia. For primary culture, the immunopanned microglia were trypsinized and subjected to culture. For RNAseq, 2 ml Trizol (Thermo, 15596018) was directly applied to the panning dish to collect the immunopanned microglia.

Alternatively, for nucleus isolation, co-IP, and chromatin interaction experiments, microglia were enriched using Percoll density centrifugation. Briefly, 10x HBSS was added to Percoll at 1:9 ratio to achieve a 100% stock isotonic Percoll (SIP) solution, which was then diluted to 70%, 37%, and 30% SIP with 1x HBSS. Brain cells isolated from three P7 mouse pups were resuspended in 4 mL 37% SIP, layered onto 4 ml 70% SIP in 15 mL falcon tube, and was then overlayed by 4 mL 30% SIP and 2 mL 1x HBSS. This percoll density was then spin at 500 g, 24 min at room temperature in a swing-bucket centrifuge, with start and brake set to zero. 2 mL of the interface between 70% and 37% SIP, in which microglia were concentrated, was collected, and diluted with 8 mL 1x HBSS, followed by centrifugation at 500 g, 7 min. Microglia could be retrieved from the pellet.

### Microglia primary culture and treatment

Serum-free primary microglia culture was performed as previously described ^70^ with minor modification. Microglia isolated by immunopanning were cultured in microglia culture medium (MGM): DMEM/F12 (Gibco, 1133032) with N2 supplement (Gibco, 17502048), GlutaMax (Gibco, 35050061), and 5 μg/mL N-Acetyl Cysteine (Sigma, A9165), supplemented with TGF-β2 (Peprotech, 100-35B, 2 ng/mL)/IL-34 (R&D, 5195-ML/CF, 100 ng/mL)/Cholesterol (Sigma, C8667, 1.5 μg/mL). Medium were changed every 2∼3 days.

For co-culture experiments, primary microglia from one P7 mouse was seeded in one 35 mm culture dish, expand for one week before subculture with neural progenitor or neuron. Microglial conditional medium was collected prior to subculture.

For lentivirus transfection, primary microglia were seeded at the density of 3 × 10^4^ cells per 12 mm coverslip in 24-well plate, cultured for 2 days before transfection. Lentiviruses were added at MOI 100, incubated for 6 h, before changed to new culture medium. Cells were fixed in 4% PFA and subjected to immunostaining staining analysis 2 days after transfection.

For drug treatments, primary microglia were seeded at the density of 1.5 × 10^4^ cells per 12 mm coverslip in 24-well plate, cultured for 5 days before treatment. Drugs and the final concentrations are: Thapsigargin (MCE, HY-13433, 1 μM), Importazole (MCE, HY-101091, 10 μM), BV02 (MCE, HY-101985, 2.5 μM), Brefeldin A (MCE, HY-16592, 10 μM) PD98059 (MCE, HY-12028, 10 μM), H89 (MCE, HY-15979A, 10 μM), Sotrastaurin (MCE, HY-10343, 10 μM), AKT inhibitor VIII (MCE, HY-10355, 5 μM), D4476 (MCE, HY-10324, 10 μM). Cells were fixed in 4% PFA and subjected to immunostaining staining analysis 24 h post treatment.

### Microglia-neural progenitor cell or neuron co-culture

Neural progenitor cell culture was performed as previously described with minor modifications ^71^. In brief, pregnant mice were sacrificed by CO_2_ overdose at the designated pregnancy stage (E12). Embryonic brains were dissected and subjected to digestion with Accutase (Gibco, A1110501) to isolate neural progenitor cells. The cells were then subjected to suspension culture at the density of 2 × 10^5^ cells/mL in neural progenitor culture medium (NPCM): DMEM/F12 with B27 supplement (Gibco, 17504044), 0.6% Glucose (Sigma, G8270), 5 mM HEPES (Sigma, H0887), 62.5 ng/mL Progesterone (Sigma, P8387), 100 nM Putrescine (Sigma, P5780), 10 μg/mL ITSS (Roche, 100053423), 1.83 μg/mL Heparin (Sigma, H0200000), supplemented with 20 ng/mL EGF (Sigma, E1257) and 5 ng/mL FGF (Sigma, F0291), and were maintained for 3 days before subculture.

For microglia-neural progenitor direct co-culture, neurosphere were dissociated using Accutase and the resulting neural progenitors seeded onto PDL-coated coverslips in NPCM at 1 × 10^5^ cells per coverslip. On the next day, microglia were subcultured, resuspended in MGM: NPCM (without EGF/FGF) (1:1) and added to coverslips with or without neural progenitors at the density of 5 × 10^4^ cells per coverslip.

For microglia-neuron co-culture, neural progenitors isolated from embryonic brain were seeded onto PDL coated coverslip at 1x 10^5^ per coverslip in NPCM, and changed to EGF/FGF-free NPCM on the next day to allow differentiation into neurons. On the 4^th^ day, microglia were subcultured, resuspended in MGM: NPCM (without EGF/FGF) (1:1) and added to coverslips with neurons at the density of 5 × 10^4^ cells per coverslip.

### Mixed glia culture

P1 mice were euthanized on ice, and brains were removed. Cerebella and meninges were discarded. The tissue was minced with a razor blade into < 1 □ mm³ fragments in dissection buffer (PBS with 33 □ mM D-glucose). Fragments were sequentially triturated using 5 □ mL, 1 □ mL, and 200 □ µL pipettes. The resulting cell suspension was centrifuged at 250 □ g for 10 □ min. Cells were resuspended in DMEM (high glucose) with 10% FBS and 1% penicillin/streptomycin, then seeded onto uncoated 10 □ cm dishes (1–2 brains per dish) in 5 □ mL medium. After 24 □ h, medium was replaced to remove debris, then changed every 2–3 □ days. At ∼90% confluence (∼7 □ days), cells were passaged using 0.25% trypsin-EDTA (2 □ min, 37 °C), centrifuged at 200–250 □ g for 3–5 □ min, and reseeded onto coverslips at 1.5 × 10 □ cells per coverslip.

### AAV and stereotypic injection

A previously reported AAV mediated gene delivery system was adopted for microglial Cx43 overexpression (10.1101/2024.07.09.602653). In brief, mouse Cx43 coding sequence was cloned and inserted into the pAAV-mIBA1-EGFP-WPRE-4 × miR9.0T-pA plasmid to replace the EGFP coding sequence. For nucleus-directed Cx43 delivery, 1 × SV40-NLS was cloned to the N-terminus of Cx43. The plasmids were then packaged with AAV11 serotype in HEK-293T cells, concentrated and exchanged into PBS containing 0.001% Pluronic F68 (Thermo, 24040032) via Amicon® Ultra centrifugal filters (Merck Millipore, UFC910024). For stereotypic injection of P2 mouse hippocampus, mouse pups were anesthetized by placing on ice for 3 min, and maintained anesthesia with isoflurane mask during injection. The skull was exposed by a 1 cm incision of skin at the midline. A 5 µL Microliter Syringe (Hamilton, 87930) was used for injection. Injection coordinates (from Lambda) were: x = 0.5, y = 0.75, z = 1.8. 0.3 μL of 1 × 10^13^ titer AAV were injected at the speed of 0.1 μL/min. The needle was left at the injection site 3 mins post-injection, and slowly retracted. The skins were re-attached with application of 5 μL 3M Vetbond^TM^ tissue adhesive.

### Lentivirus treatment

Lentivirus was used to transfect primary microglia culture. For overexpression of Cx43 NLS-GFP fusion proteins, the coding sequence of putative NLSs of Cx43 (237-290, and 343-373) was cloned to the pcSLenti-CMV-EGFP-3xFLAG-PGK-puro-WPRE3, at the 3’ end of GFP. For overexpression of Cx43 NLS deleted mutants, wild type and NLS-deleted mutant Cx43 was cloned to the pcSLenti-CMV-EGFP-3xFLAG-PGK-puro-WPRE3. The expression vectors and the backbone plasmid (ObiO) were transfected into 293FT cells using transfection reagent (ObiO, OGTR(C)20131002) for virus packaging. Lentivirus were collected 60 h post transfection.

### Immunofluorescence staining

Mice were anesthetized by the i.p. injection of 20% urethane at 10 μL/g, followed by transcardial perfusion of PBS and 4% paraformaldehyde (PFA). The brains were then isolated and subjected to post-fixation with 4% PFA at 4 □ overnight, then immersed in 30% Sucrose for cryoprotection, before cryostat sectioning for 20 μm thick slices. For cell culture, the cells on coverslips were rinsed once with ice-cold PBS, and fixed with 4% PFA at room temperature for 10 min, then rinsed trice with PBS. Brain slices or coverslips were permeabilized with 0.5% Triton-X100 in PBS (0.5% PBST) for 1 hour, blocked with 2% BSA in PBST for 1 hour, and incubated with primary antibody overnight at 4 □, followed by secondary antibody, counterstained with DAPI, and mounted for microscopic observation. Primary antibodies and dilution are listed below: Rabbit-anti-Cx43 (Merck, C6219, 1:200), Goat-anti-IBA1 (Abcam, ab5076, 1:400), Rat-anti-CD11b (Abcam, ab8878, 1:200), Goat-anti-GFAP (Abcam, ab53554, 1:400), Rabbit-anti-Aldh1l1 (Abcam, ab87117, 1:200), Rat-anti-MBP (Millipore, MAB386, 1:200), Goat-anti-PDGFRa (R&D, AF1062, 1:400), Rabbit-anti-Olig2 (Millipore, AB9610, 1:200), Rat-anti-Nestin (Oasis biofarm, OB-PRT109), Rabbit-anti-NeuN (Abcam, ab177487, 1:200), Rabbit-anti-DCX (Abcam, ab18723, 1:200), Guinea pig-anti-DCX (Oasis biofarm, OB-PGP025, 1:200), Rabbit-anti-cleaved caspase3 (CST, 9661, 1:200), Rabbit-anti-phospho-MLKL (CST, 37333, 1:200), Rabbit-anti-cleaved GSDMD (Abcam, ab255603, 1:200), Rabbit-anti-Ki67 (Thermo, MA514520, 1:200), Rat-anti-Dectin-1(CLEC7A) (Invivogen, mabg-mdect-2, 1:200), Rat-anti-CD16/32 (FCGR3) (Biolegend, 101320, 1:100), Rabbit-anti-MEF2C (CST, 5030, 1:200), Mouse-anti-C1q (Hycult Biotech, HM1096BT, 1:200), Guinea pig-anti-Vglut1 (Synaptic Systems, 135304, 1:200), Mouse-anti-Vgat (Synaptic Systems, 131011, 1:200), Rat-anti-HA (Merck, 11867423001, 1:200), Goat-anti-GFP (Abcam, ab5450, 1:200), Rabbit-anti-phospho-AKT (S473) (CST, 4060, 1:200).

Tunel labelling was performed using In Situ Cell Death Detection Kit, TMR red (Roche, 12156792910), according to the manufacturer’s protocol. The immunostaining procedures were performed after Tunel labelling.

Images were captured using a VS200 slide scanner, FV3000 confocal microscope, or SpinSR spinning disk confocal microscope (Olympus). Super-resolution images were captured using a SpinSR spinning disk confocal microscope.

### Immunoelectron microscopy

Following post-fixation for 2 □ h in 4% paraformaldehyde (PFA) at 4 °C, the brains were washed with phosphate-buffered saline (PBS: 50 mM, pH 7.4); and sectioned coronally with a vibratome (VT1200S, Leica Biosystems) at 50 µm of thickness, and stored in cryoprotectant solution (30% ethylene glycol, 30% glycerol in PBS) at -20 □ °C until use.

Brain sections containing the hippocampus *stratum lacunosum moleculare* (SLM) from a P7 C57BL/6J female mouse (Jackson laboratory) (Bregma –1.55 mm to -2.03 mm) ^72^ were selected for further processing. The selected sections were quenched in 0.3% H_2_O_2_ (Fisher Scientific, cat# 202762) in PBS for 5 min. Afterward, the sections were incubated in 0.1% NaBH_4_ in PBS for 30 min, followed by 3 times 10-min washes in PBS. Brain sections were then incubated in a blocking buffer solution containing 10% fetal bovine serum (Jackson ImmunoResearch Labs, cat# 005-000-121), 3% bovine serum albumin (Sigma-Aldrich, cat# 9048-46-8), and 0.03% Triton X-100 in TBS for 1 h at room temperature (RT). They were then incubated overnight in blocking buffer solution with the primary anti-connexin-43 antibody produced in rabbit (1:150; Sigma-Aldrich, cat# C6219) at 4 °C. The following day, the brain sections were washed with Tris-buffered saline (TBS; 50 mM, pH 7.4), then incubated with a 1.4 nm Nanogold®-Streptavidin antibody (1:100; Avantor, cat# 414004-023) in TBS for 2 h at RT. Then, sections were washed by ddH_2_O (2%) for 3 10-min washes. The staining was revealed with HQ Silver enhancement kit (Nanoprobes, cat# #2012-45ML).

Brain sections were incubated in 3% potassium ferrocyanide (in phosphate buffer; BioShop, cat# PFC232.250) combined (1:1) with 4% aqueous osmium tetroxide (cat# 19170, Electron Microscopy Sciences, Hatfield, PA, United States) for 1 h, washed in phosphate buffer (PB; 100 □ mM, pH 7.4), incubated in 1% thiocarbohydrazide (in double distilled water (ddH_2_O); Electron Microscopy Sciences, cat# 2231-57-4) for 20 □ min, washed in ddH_2_O, incubated in 2% osmium tetroxide (in ddH_2_O), then dehydrated in ascending concentration of ethanol (2 times in 35%, 50%, 70%, 80%, 90%, and 3 times in 100%) followed by 3 times incubation in propylene oxide. Post-fixed sections were embedded in Durcupan ACM resin (MilliporeSigma, cat# 44611–44614) for 24 □ h, placed between two ACLAR® embedding sheets (Electron Microscopy Sciences, cat# 50425-25), and resin was polymerized at 55 □ °C for 72 □ h. The hippocampus SLM was extracted, glued on a resin block, and cut into 70 nm thick ultrathin sections using a Leica ARTOS 3D ultramicrotome (Leica Biosystems).

Ultrathin sliced sections were mounted on a dust-free silicon nitride chip, glued on specimen mounts, and inserted into a Zeiss Crossbeam 350 focused ion beam scanning electron microscope (FIB-SEM). The region of interest, the hippocampus SLM, was identified using backscattered electron (BSE) and secondary electron (SE2) detectors at 10 kV for initial focus, then at 1.4 kV when the samples were at an appropriate working distance for high-resolution imaging of brain ultrastructure. These steps were controlled using SmartSEM software (Fibics). Microglial cell bodies, identified by their overall shape, unique heterochromatin pattern pockets, as well as their long and narrow stretches of endoplasmic reticulum ^73^. Immunogold labeling was recognized on microglia cells as discrete electron-dense puncta, and a matched negative control (omission of primary antibody) was included to assess nonspecific nanogold background. The connexin-43-positive microglia were imaged at 5 nm of resolution, and the images were exported with the TIFF file format using the Zeiss ATLAS Engine 5 software (Fibics).

### Acute brain slice preparation and related experiments

The acute brain slices were prepared as previously described. Briefly, mice were anesthetized by i.p. injection of 2.5% Avertin at 200 μL/10 g and euthanized by rapid decapitation with sharpened scissors. The brains were rapidly isolated, stationed at the specimen holder, immersed in the ice-cold Sucrose-based artificial cerebrospinal fluid (ACSF) with constant carbonation (95% O_2_, 5% CO_2_), and subjected to vibratome sectioning to obtain 300 μm slices. Slices were then recovered in ACSF at 32 □ for 30 min, and transferred to the carbonated ACSF at room temperature and recovered for another 30 min before experiments.

### Dye uptake experiment

The dye uptake experiment is a widely used assay to monitor the opening of hemichannels; this assay utilizes poor selectivity of connexin hemichannels, which allow passage of small molecules with MW < 1.5 kDa. Several dyes were used for dye uptake experiments, including EtBr, DAPI, carboxyfluorescein, etc., all of them cannot cross the lipid bilayer but permeate through connexin hemichannels. EtBr dye was selected over other dyes such as carboxyfluorescein and DAPI due to (i) carboxyfluorescein would diffuse across the cell body after entering through hemichannel, making it implausible to perform cell type specific dye uptake analysis; (ii) while both EtBr and DAPI would bind to DNA after entering the cell, which allow cell type specific analysis, the autofluorescence of Aβ plaque interferes with DAPI signal. Dye uptake experiment was performed on acute brain slices as previously described. Briefly, acute brain slices were transferred onto 70 μm cell strainers put in the 6-well plate filled with the recording ACSF. Alternatively, CBX (200 µM), and TAT-Cx43_266-283_ (50 μM) were added to the recording ACSF. After 15 min incubation, EtBr dye was added to the recording buffer at 5 µM, and incubated for 10 min. The slices were then rinsed 3 times with the recording ACSF (5 min each), fixed with 4% PFA for 1 hour before immunostaining.

For the dye uptake experiment in primary microglial culture, cells were rinsed twice with Hank’s balanced salt solution (HBSS) (Ca^2+^/Mg^2+^ free), and preincubated in HBSS with or without TAT-Cx43_266-283_ for 15 min. As a negative control, CaCl_2_ was added to the HBSS at 1.8 mM. EtBr was then added to 5 μM, and further incubated with cells for 10 min. Cells were then fixed with 4% PFA for 10 min and subjected to microscopic analysis.

### RNA-sequencing

RNA-sequencing was performed on acute-isolated microglia from P7 Cx43^mglcKO^ mice and littermate controls. RNA was extracted with a RNEasy Plus mini-isolation kit (Qiagen), and the RNA quality was examined by electrophoresis. The sequencing was performed using the DNBSEQ platform. The raw data were filtered with FastQC to remove reads of low quality, reads with adaptor sequences, and reads with high levels of N bases, ranging from 98.65 to 99.96. The filtered reads were mapped to the reference Genome (*Mus musculus*, GCF_000001635.26_GRCm38.p6) using the HISAT-Bowtie2 pipeline. The average mapping ratio with reference genome is 98.74%, the average mapping ratio with gene is 81.80%; 18571 genes were identified. Differential expression analysis was performed using DESeq2. Pair-wise comparisons were performed between them. Differentially expressed genes were determined by a threshold of FC > 1.5, p < 0.05, and subjected to pathway enrichment analysis using DAVID (https://david.ncifcrf.gov/).

### Nucleus isolation

Microglia and brain cells were resuspended in 1 mL homogenization buffer (HB; 0.25 M sucrose, 25 mM KCl, 5 mM MgCl_2_, 1 mM dithiothreitol (DTT), 50 mM Tris-HCl pH 7.4, supplemented with protease inhibitor cocktail (Beyotime Biotech)). 50 μL cell suspension was collected as whole cell lysate input. Triton-X100 was added to the rest of cell suspension at 1% and triturated 10 times to disrupt the plasma membrane. The resulting suspension was added to a sucrose cushion (1.8 M sucrose, 5 mM MgSO4, 0.1 mM EDTA, 10 mM Tris-HCl pH 7.4), and centrifuged for 80 min at 21,100 g. The upper layer was collected as cytoplasmic fraction, and the pellet was resuspended for centrifugation at 4,000 g for 10 min. The nuclear fraction was then supplemented with NaCl (300 mM), Triton-X100 (1%), and SDS (0.2%), and sonicated (40% amplitude, 2 sec on/off for 10 cycles) for sufficient tissue lysis. The protein concentration was then determined by BCA analysis.

### Cx43 multimer analysis

To examine the multimeric state of Cx43, the cytosolic fraction and nucleus fraction of microglia with 1% Triton-X100 and 4 μg/mL benzonase (Yeasen), and sonicated (40% amplitude, 2 sec on/off for 10 cycles). The lysate was divided into four equal aliquots, subjected to naïve, DTT-only, DSP-only, DSP+DTT treatment, respectively. For DSP crosslinking, the lysates were treated with 100 μg/mL Pierce™ DSP (Thermo, 22586) on ice for 30 min, and quenched with glycine stock (20 μL/mL, pH9.2), on ice for 30 min. The lysate was then supplemented with 1 M glycine (pH 7.2) at 10 μL/mL to lower the pH and with 150 mM NaCl. Cx43 was then immunoprecipitated by rabbit-anti-Cx43 antibody diluted at 1:300 (Merck, C6219) at 4 °C overnight, and then Protein A/G agarose beads (Santa Cruz, sc-2003) for 2 h at room temperature. The beads were washed once with wash buffer (50 mM Tris-HCl pH 7.4, 150 mM NaCl, 0.5 mM NP40, 1% Triton-X100), and twice with wash buffer supplemented with 0.1% SDS. The beads were then eluted by incubation with 4x loading buffer (0.2 M Tris-HCl pH 6.5, 8% SDS, 4 mg/ml bromophenol blue, 32% glycerol) with or without 0.4 M DTT, at 95°C for 3 min. The proteins eluted were then loaded onto a PAGE gel and incubated with goat anti-Cx43 antibody (LSBio, LS □ B9771).

### Western blot

Protein lysates were prepared using RIPA buffer (50 mM Tris-HCl, pH 7.4, 150 mM NaCl, 1% NP-40, 0.5% sodium deoxycholate, 0.1% SDS) supplemented with protease and phosphatase inhibitors. Protein concentrations were determined using a BCA assay. Laemmli Sample buffer was added to the protein lysis, and incubated at 95 °C for 5 min. Proteins were separated by SDS-PAGE on polyacrylamide gels and transferred onto PVDF membranes. Membranes were blocked for 1 h at room temperature in 5% non-fat dry milk or 5% BSA in Tris-buffered saline containing 0.1% Tween-20 (TBST). Blots were incubated overnight at 4 °C with primary antibodies. After washing with TBST, membranes were incubated with HRP-conjugated secondary antibodies for 1 hour at room temperature. Protein bands were visualized using enhanced chemiluminescence (ECL) substrate and imaged using a chemiluminescence detection system. Primary antibody used were: Rabbit-anti-Cx43 (Merck, C6219, 1:2000), Goat-anti-Cx43 antibody (LSBio, LS □ B9771, 1:2000), mouse-anti-LMNB1 (Abcam, ab324446, 1:1000), Rabbit-anti-H3 (Abclonal, A2348, 1:1000), Mouse-anti-GAPDH (Proteintech, 60004-1-Ig, 1:1000), Mouse-anti-NDUFA9 (Invitrogen, 459100, 1:1000), Rabbit-anti-FACL4 (ACSL4) (Abcam, ab155282, 1:1000), Goat-anti-IBA1 (Abcam, ab5076, 1:1000), Rabbit-anti-Aldh1l1 (Abcam, ab87117, 1:1000), Mouse-anti-GFP (Abclonal, AE012, 1:1000), Rabbit-anti-phospho-AKT (S473) (CST, 4060, 1:1000), Rabbit-anti-pan AKT (CST, 4691, 1:1000), Rabbit-anti-phospho-Cx43 (S373) (Affinity, AF8264, 1:1000).

### ChIP

Acute-isolated microglia by percoll method (1 × 10 □ per sample) were crosslinked with 1% formaldehyde for 10 □ min at room temperature, and the reaction was quenched with 125 □ mM glycine. Cells were then lysed in ChIP Lysis Buffer. Chromatin was sonicated to an average fragment size of 200-1000 □ bp, and debris was removed by centrifugation. For each immunoprecipitation, 25 μg of DNA was diluted 1:10 in RIPA Buffer and incubated with the primary antibody (Cx43 antibody, Sigma, C6219, 1:200) for 1 □ h at 4 □ °C. Blocked Magna ChIP Protein □ A/G beads (Sigma, 16-663) were added and incubated overnight at 4 □ °C. Beads were washed sequentially with low-salt, high-salt, and LiCl wash buffers. Bound complexes were eluted with Elution Buffer, and crosslinks were reversed overnight at 65 □ °C with RNase □ A, followed by proteinase □ K treatment. Purified DNA was used to prepare libraries, which were sequenced on an Illumina platform.

### Cut&Tag assay

Concanavalin A-coated magnetic beads were prepared according to the manufacturer’s instructions and incubated with the cells to immobilize them. After removal of unbound beads, cells were permeabilized with digitonin to allow antibody access. Samples were then incubated overnight at 4 °C with a primary antibody (Cx43 antibody, Sigma, C6219, 1:100), followed by incubation with a secondary antibody. A pA/G-Tn5 transposase fusion protein was added to bind the antibodies, and Mg² □ was introduced to activate the Tn5 transposase, which simultaneously fragments the DNA and adds sequencing adapters. Following tagmentation, DNA was purified, and libraries were amplified by PCR. Libraries were sequenced on an Illumina platform.

### Cut&Run assay

Concanavalin A-coated magnetic beads (10 □ µL slurry) were activated in Binding Buffer. Cells (1 × 10 □ per sample) were harvested, washed twice in Wash Buffer, and resuspended in 100 □ µL Wash Buffer. Activated beads were added and incubated for 15 □ min at room temperature. Bead-bound cells were resuspended in Dig-Wash Buffer containing primary antibody (Cx43 antibody, Sigma, C6219, 1:100) at 4 □ °C overnight, followed by PA/G-MNase fusion protein for 1 □ h at 4 □ °C, then washed again. Chromatin digestion was initiated by adding 2 □ µL of 100 □ mM CaCl □ and incubating for 30 □ min at 4 □ °C on a rotator. The reaction was stopped with 2X STOP Buffer (30 □ min at 37 □ °C). Released chromatin fragments were collected by magnetic clearance and purified by phenol-chloroform extraction. Libraries were prepared using the QIAseq Ultralow Input Library Kit and sequenced as 150 □ bp paired-end reads on an Illumina platform.

### Co-IP

For Cx43 co-IP from cell fractions including the nucleus, microglial cell lysis was performed as previously reported ^74^. Briefly, microglia were collected and resuspended in HEPES-Glycerol-NaCl buffer (HGN buffer: 10 mM HEPES-NaOH, pH 7.4, 10% Glycerol (v/v), 165 mM NaCl, with protease cocktail). A 4-step sequential lysis were performed, each for 1 hour at 4 □, followed by spin at 8000 g, 10 min to separate soluble and insoluble fractions. Lysis buffer 1 (LSB1) was with HGN buffer supplemented with 0.1% Triton X-100, 0.5 mM MgCl2, 0.5 mM DTT, 0.5 unit/μL Benzonase; Lysis buffer 2 was with LSB1 supplemented with 0.2% Triton X-100; Lysis buffer 3 was with LSB1 supplemented with 50 mM NaCl; Lysis buffer 4 was with LSB1 supplemented with 100 mM NaCl. The supernatant was pooled together and proceed with Cx43 co-IP (Merck, C6219, 1:300), with IgG as control, by overnight incubation at 4 °C, and then with Protein A/G agarose beads (Santa Cruz, sc-2003) for 2 h at room temperature. The beads were washed 4 times with HGN buffer, and eluted with SDS sampling buffer.

### Mass spectrometry

IP samples were separated on 12% PAGE gel, and stained with Coomassie Brilliant Blue R-250. Each protein sample was taken, and the volume was made up to 100 μL with DB lysis buffer (6 M Urea, 100 mM TEAB, pH 8.5), trypsin and 100 mM TEAB buffer were added, sample was mixed and digested at 37 °C for 4 h. Formic acid was mixed with digested sample, adjusted pH under 3, and centrifuged at 12,000 g for 5 min at room temperature. The supernatant was slowly loaded to the C18 desalting column, washed with washing buffer (0.1% formic acid, 3% acetonitrile) 3 times, then added elution buffer (0.1% formic acid, 70% acetonitrile). The eluents of each sample were collected and lyophilized. Prepare mobile phase A (99.9% water, 0.1% formic acid) and B (80% acetonitrile, 0.1% formic acid). The lyophilized powder was dissolved using 10 µ LA solution, centrifuged at 14,000g for 20 min at 4 °C, and 200 ng of the supernatant sample was injected into the sample for liquid-quality detection. A Vanquish Neo upgraded UHPLC system was used with a C18 pre-column of 174500 (5 mm × 300 μm, 5 μm, thermo) heated at 50 °C in a column oven, and a C18 analytical column of ES906 (PepMap TM Neo UHPLC 150 µm × 15 cm, 2 μm, thermo). A Thermo orbitrap astral mass spectrometer mass spectrometer was used with an Easy-spray (ESI) ion source. The raw files were searched and analyzed using the DIA-NN library search software, according to the Mus_musculus_uniprot_2025_05_15_Swissprot.fasta protein database. The mass deviation of precursor ions and fragment ions is automatically detected and corrected. The fixed modification is C carbamidomethylation, while N-term M excision is considered a variable modification. Up to 2 missed cleavage sites are allowed. To improve the quality of analysis results, the DIA-NN software further filters the search results, retaining only peptides with a Global.Q.Value < 0.01 and proteins with a PG.Q.Value < 0.01. A protein with a fold change greater than or less than a certain value was defined as a differentially expressed protein.

### Statistical analysis

Statistical significance between groups was determined with GraphPad Prism software. The unpaired two-sided t-test was used to determine the difference between two groups with normal distribution, the Mann-Whitney test was used to determine the difference between two groups with non-normal distribution. One-way analysis of variance (ANOVA) or Two-way ANOVA was used to determine the difference when there were more than 2 groups where applies. The Pearson correlation test was used to determine the correlation between variables. Data distribution was tested by Shapiro–Wilk test. A probability of *p* < 0.05 was considered statistically significant. All significant statistical results are indicated within the figures with the following conventions: * *p* < 0.05, ** *p* < 0.01, *** *p* < 0.001, **** *p* < 0.0001. Error bars represent ± standard error of the mean (SEM). No statistical methods were used to predetermine sample sizes. The sample size per group was determined from previous publications using a similar methodology. One mouse from each litter was randomly selected for each experiment. Investigators were blinded to group allocation during data analysis. All in vitro experiments were performed at least three times.

### Data availability

RNA-seq data have been deposited in the NCBI GEO database under the accession number GSE298333. The mass spectrometry proteomics data have been deposited to the ProteomeXchange Consortium (http://proteomecentral.proteomexchange.org) via the iProX partner repository ^75^ with the dataset identifier ProteomeXchange: PXD078951.

## Acknowledgements

This work was supported by Shenzhen Fundamental Research Program (RCJC20231211090018040 and ZDSYS20220606100801003 to C.Y.), National Natural Science Foundation of China (W2511095 and 32271034 to J.N., 32300791 to Y.S.), Shenzhen Medical Research Fund (A2303014 to Y.S.), Guangdong Basic and Applied Basic Research Foundation (2026B1515020011, 2023A1515010651 to Y.S.)

## Author contributions

C.Y., A.V., J.N., and Y.S. conceptualized the study. C.Y., J.N., and A.V. supervised the study and ensured the coordination and resource support of the project. Y.S. performed most of the experimental work, analysis, and visualization. Q.F. performed the biochemistry experiments. Q.W. and S.Z. performed the bioinformatics analysis.

P.K. performed the immunoelectron microscopic analysis under supervision from M-E.T., Q.F., H.L., C.W., X.C., Z.W. provided assistance with animal breeding and behavioral tests. R.F. acquired the human sample. Y.S. wrote the original manuscript. C.Y., A.V., J.N., and H.C. revised the manuscript.

## Competing interests

The authors declare no competing interests.

## Supplementary figures

**Fig. S1.**
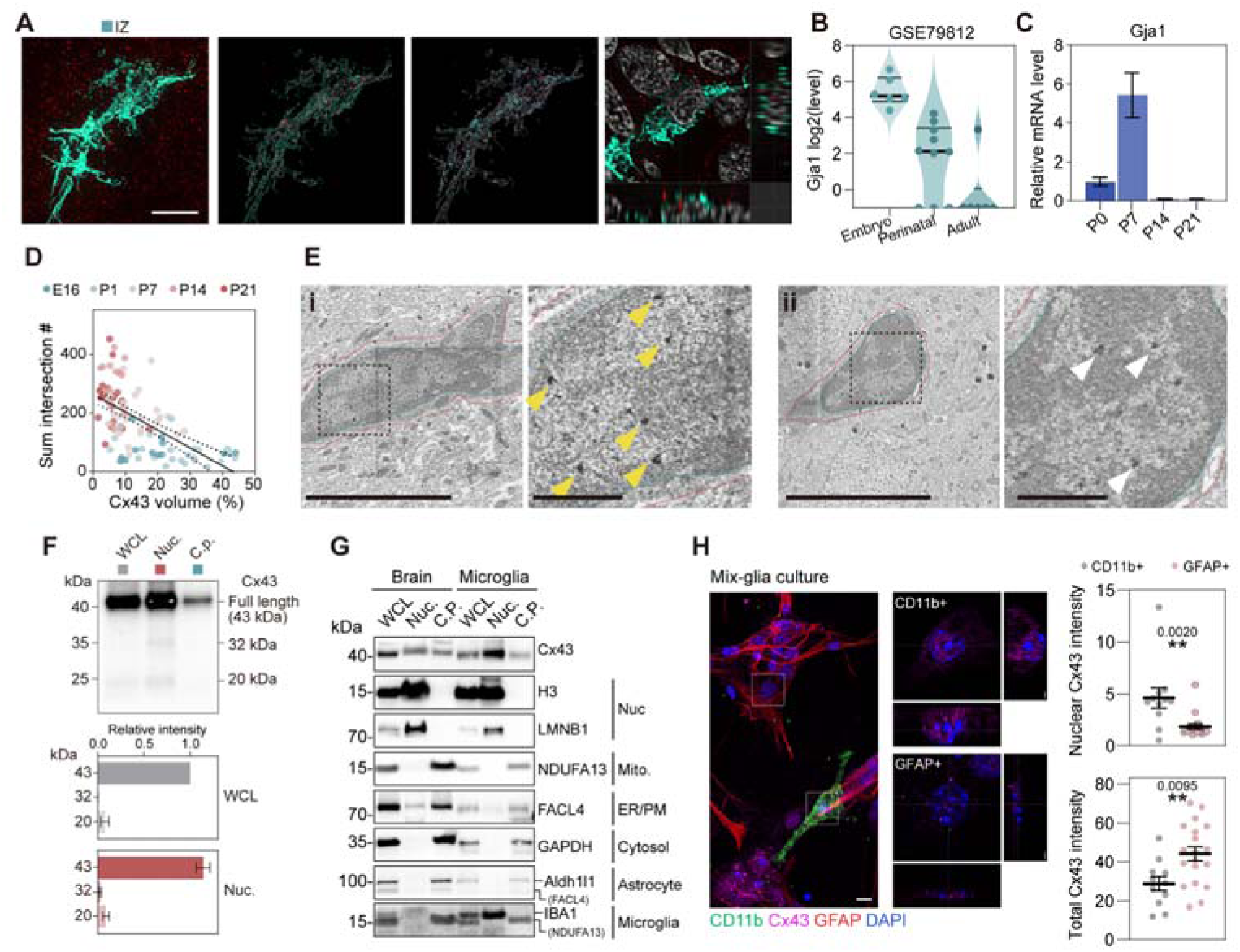
Developmental profiling of microglial Cx43 expression and function. **(A)** Representative superresolution images of IBA1^+^ microglia from intermediate zone (IZ), together with the 3D reconstructions of IBA1/DAPI masked Cx43 signal, and the orthogonal views (Scale bar, 10 μm). **(B)** Cx43 (*Gja1*) expression in mouse microglia in different stages, from single cell RNA-seq dataset. Plot shows median and quartile. **(C)** Cx43 (*Gja1*) expression in mouse microglia in different stages determined by RT-qPCR, normalized to Gapdh. N = 2 technical repeats. **(D)** Correlation test between Cx43 volume and microglia process sum intersection from Sholl analysis. Plot shows linear regression and 95% confidence. **(E)** Representative immunoelectron microscopy images in dorsal hippocampus, *stratum lacunosum moleculare* from a P7 wild type mice, immunogold-labeling for Cx43 (**i**) or secondary antibody-only control (**ii**). Red line, microglia cell body. Green line, microglia nucleus. Yellow arrowhead, labelled Cx43. White arrowhead, non-specific labeling in secondary antibody-only control (Scale bar, 5 μm and 1μm). **(F)** Western blot analysis of in microglia fractionation experiment. The expanded figure of Cx43 immunoblotting in Fig. 1K, showing low molecular weight bands. The putative full length, 32 kDa, and 20 kDa Cx43 band were quantified. **(G)** Western blot analysis of fractionation of whole brain cells and microglia. **(H)** Representative images of mix-glia culture, staining for Cx43, CD11b and GFAP. The Cx43 intensity in the nucleus and in the whole cell was quantified for CD11b^+^ microglia and GFAP^+^ astrocytes. N = 11 and 19 cells from 3 experiments. Results are expressed as mean ± SEM. \**p* < 0.05, \*\**p* < 0.01, \*\*\**p* < 0.001, \*\*\*\**p* < 0.0001.

**Fig. S2.**
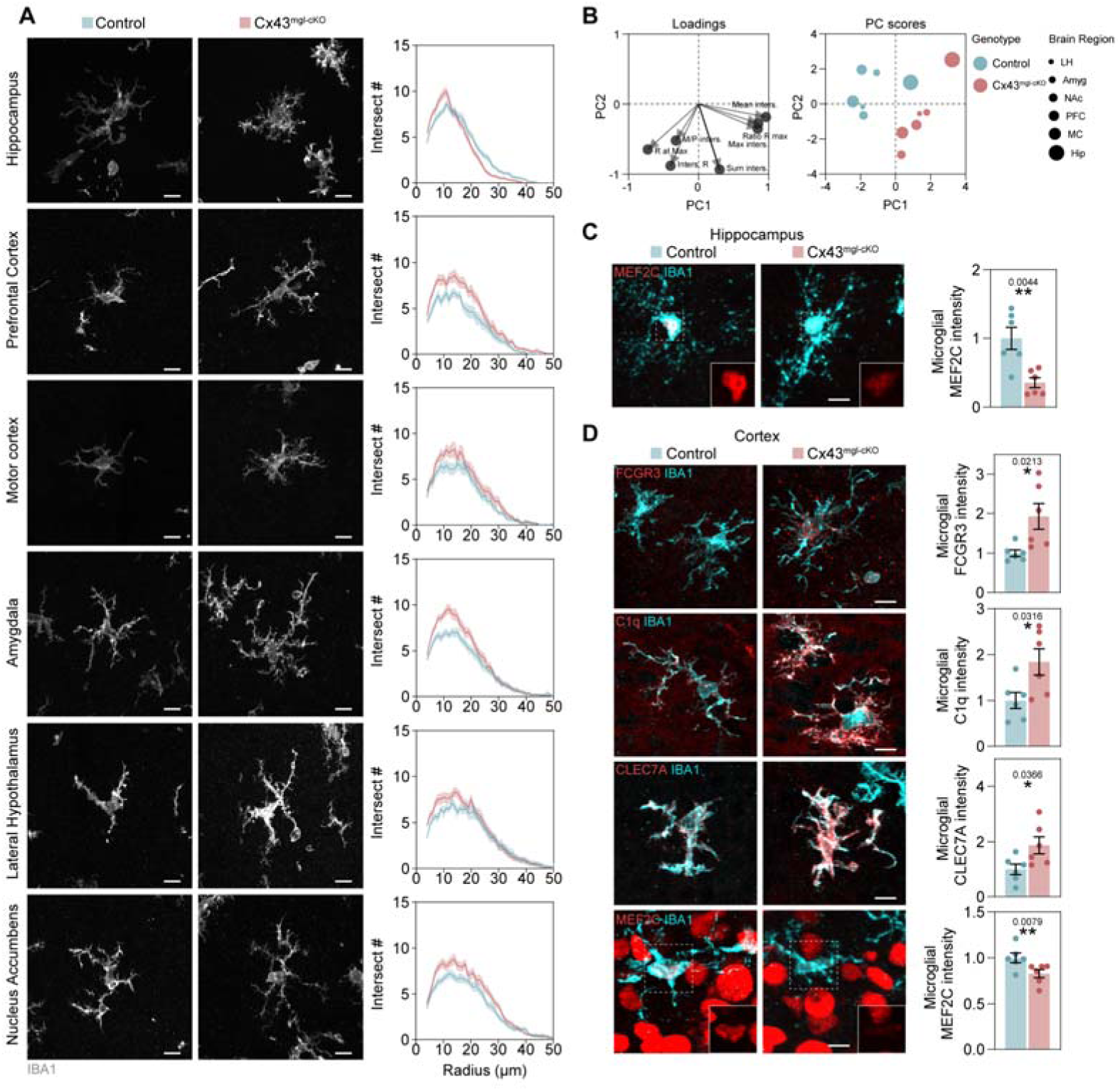
Postanatal ablation of microglial Cx43 drives neurotoxic microglia phenotype. **(A)** Representative images of P7 control and Cx43^mgl-cKO^ mouse IBA1^+^ microglia in different brain regions (Scale bar, 10 μm). Sholl analysis results are blotted. **(B)** PCA analysis of microglia morphological features from Sholl analysis. Color decodes genotype, size decodes brain regions. **(C)** Representative images of P7 control and Cx43^mgl-cKO^ mouse IBA1^+^ microglia in hippocampus, co-staining for MEF2C (Scale bar, 10 μm). Plot shows quantification of microglial MEF2C intensity. N = 6 mice. **(D)** Representative images of P7 control and Cx43^mgl-cKO^ mouse IBA1^+^ microglia in cortex, co-staining for FCGR3, C1q, CLEC7A, and MEF2C (Scale bar, 10 μm). Plots show quantification of microglial FCGR3, C1q, CLEC7A, and MEF2C intensity. N = 6 mice. Results are expressed as mean ± SEM. \**p* < 0.05, \*\**p* < 0.01, \*\*\**p* < 0.001, \*\*\*\**p* < 0.0001.

**Fig. S3.**
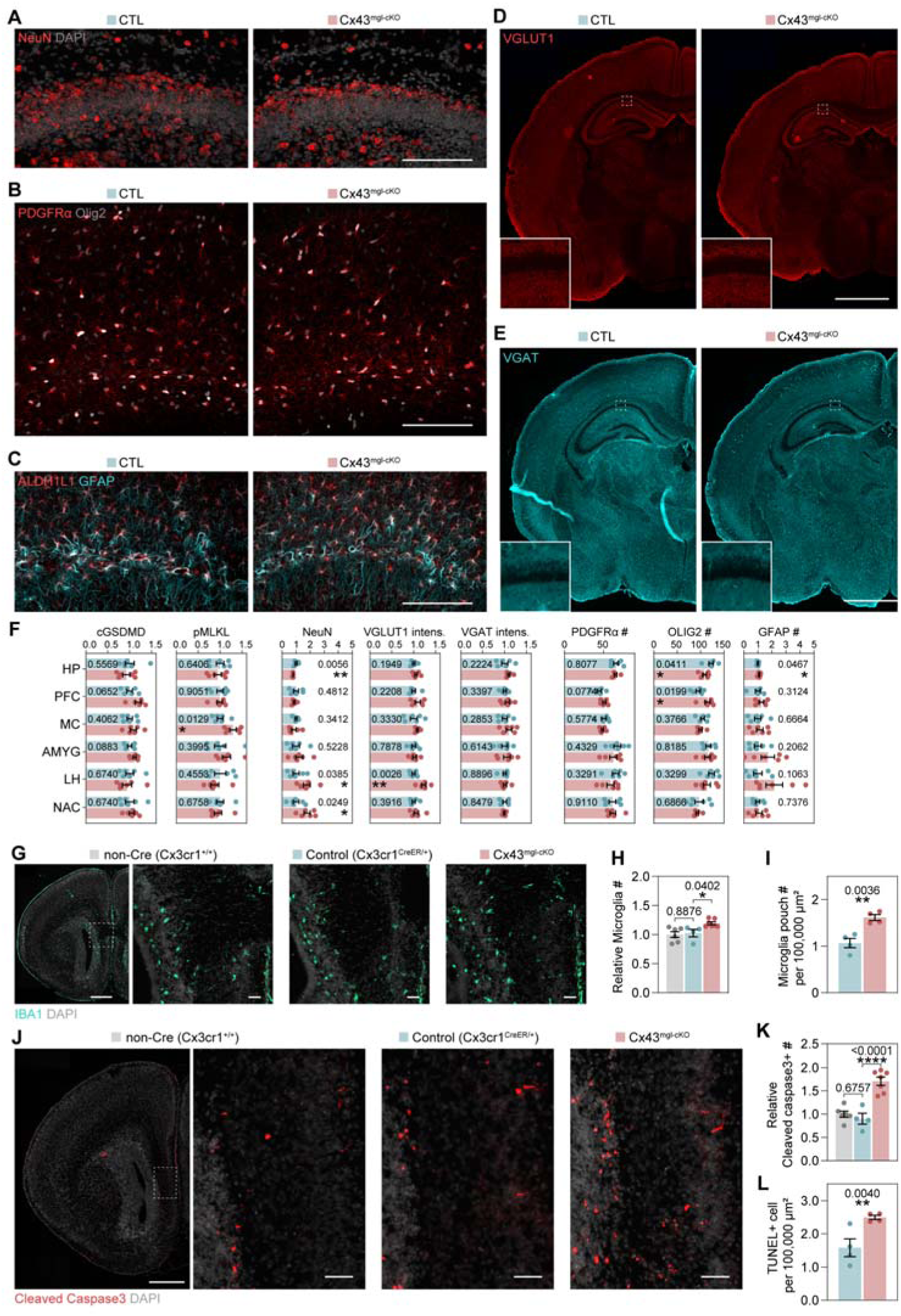
Embyronic and Postnatal deletion of microglial Cx43 promotes neuronal death. **(A-C)** Representative images of P7 control and Cx43^mgl-cKO^ mouse hippocampus, staining for NeuN (A), PDGFRα and Olig2 (B), ALDH1L1 and GFAP (C) (Scale bar, 100 μm). **(D-E)** Representative images of P7 control and Cx43^mgl-cKO^ mouse hippocampus, staining for VGLUT1 (D) and VGAT (E) (Scale bar, 500 μm). **(F)** Quantification of cGSDMD intensity, pMLKL intensity, NeuN^+^ cell number, VGLUT intensity, VGAT intensity, PDGFRα^+^, Olig2^+^, and GFAP^+^ cell number, in different brain regions. N = 6 mice. **(G)** Representative images of E18 non-Cre, control (Cx3cr1^CreER/+^), and Cx43^mgl-cKO^ mouse brain, staining for IBA1 (Scale bar, 500 and 50 μm). **(H)** Quantification of microglia number of the whole brain section. N = 6, 4, 7 mice. **(I)** Quantification of microglial phagocytic pouch number in E18 control (Cx3cr1^CreER/+^) and Cx43^mgl-cKO^ mouse brain. N = 4 mice. **(L)** Representative fluorescence microscopy images of E18 non-Cre, control (Cx3cr1^Cre/+^), and Cx43^mgl-cKO^ mouse brain, staining for cleaved Caspase 3 (Scale bar, 500 and 50 μm). **(M)** Quantification of cleaved Caspase 3+ cell number. N = 6, 4, 7 mice. **(L)** Quantification of TUNEL+ cell number in control (Cx3cr1^Cre/+^) and Cx43^mgl-cKO^ mouse brain. N = 4 mice. Results are expressed as mean ± SEM. \**p* < 0.05, \*\**p* < 0.01.

**Fig. S4.**
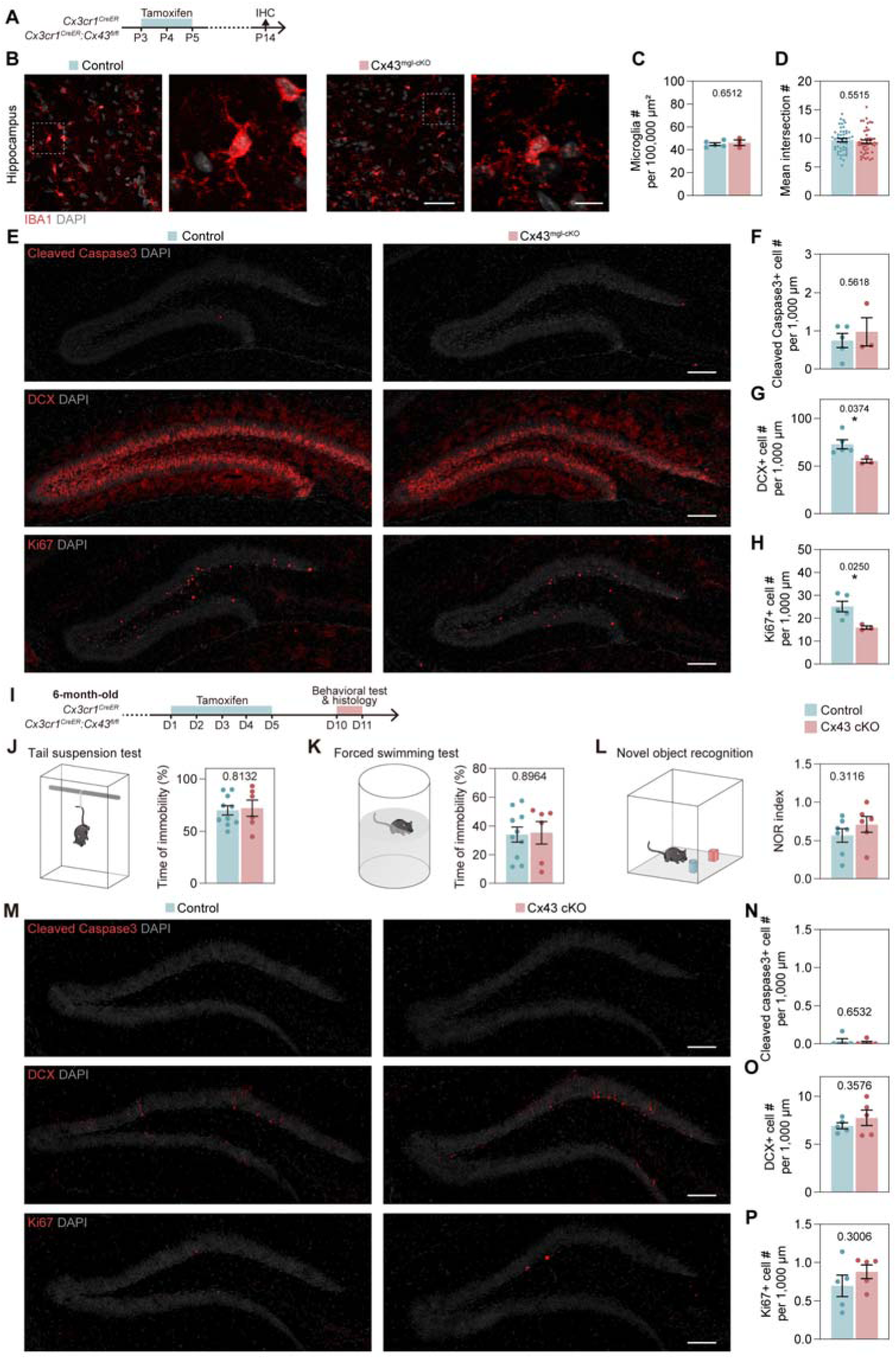
Postnatal but not adult ablation of microglial Cx43 ablation caused exhaustion of neurogenic pool and behavioral deficits. **(A)** Schematic of Cx43 conditional knockout experiment and examination at P14. **(B)** Representative images of P14 control and Cx43^mgl-cKO^ mouse hippocampus, staining for IBA1 (Scale bar, 50 and 10 μm). **(C)** Quantification of microglia number in hippocampus. N = 4 and 3 mice. **(D)** Quantification of microglia mean intersection number (Sholl analysis). N = 48 cells from 4 and 3 mice. **(E)** Representative fluorescence microscopy images of P14 control and Cx43^mgl-cKO^ mouse hippocampus, staining for cleaved Caspase 3, DCX, and Ki67 (Scale bar, 100 μm). **(F)** Quantification of cleaved Caspase 3^+^ cell number in dentate gyrus. N = 4 and 3 mice. **(G)** Quantification of DCX^+^ cell number in dentate gyrus. N = 4 and 3 mice. **(H)** Quantification of Ki67^+^ cell number in dentate gyrus. N = 4 and 3 mice. **(I)** Schematic of Cx43 conditional knockout induction in adult mice. **(J)** Quantification of immobility time in tail suspension test. N = 10, 6 mice. **(K)** Quantification of immobility time in forced swimming test. N = 10, 6 mice. **(L)** Quantification of NORT index in novel object recognition test. N = 7, 6 mice. **(M)** Representative images of adult-induced control and Cx43^mgl-cKO^ mouse hippocampus, staining for cleaved Caspase 3, DCX, and Ki67 (Scale bar, 100 μm). **(N)** Quantification of cleaved Caspase 3^+^ cell number in dentate gyrus. N = 5 mice. **(O)** Quantification of DCX^+^ cell number in dentate gyrus. N = 5 mice. **(P)** Quantification of Ki67^+^ cell number in dentate gyrus. N = 5 mice. Results are expressed as mean ± SEM. \**p* < 0.05.

**Fig. S5.**
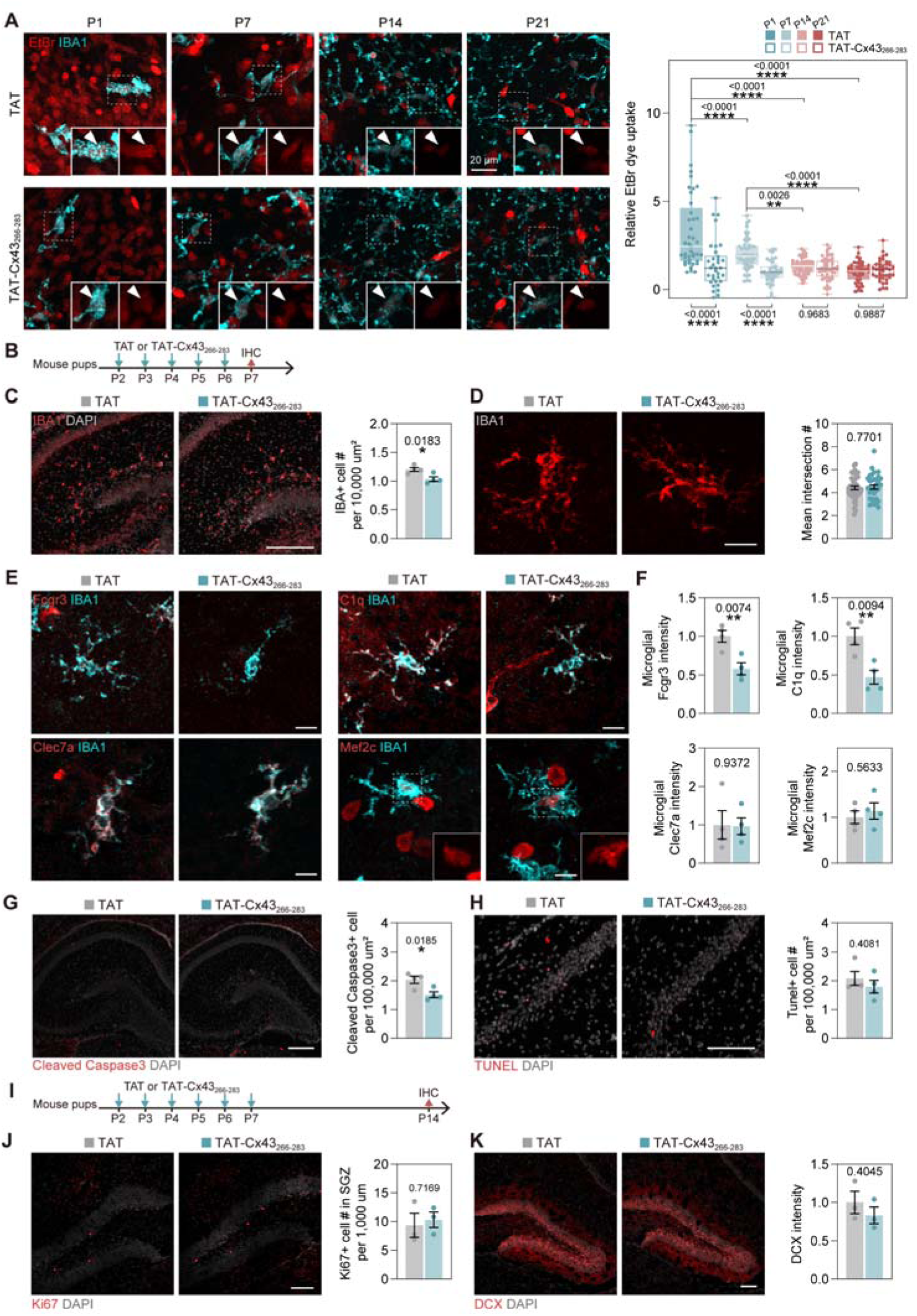
Microglial Cx43 hemichannel did not confer neuroprotective role in development. **(A)** Representative fluorescence microscopy images of EtBr dye uptake experiment performed on P1, P7, P14, P21 mice brain slices, counter-staining for IBA1 (Scale bar, 20 μm). The EtBr intensity was quantified. Plot shows median, quartile, and min/max. N = 41, 34 (P1), 53, 40 (P7), 53, 50 (P14), 42, 41(P21). **(B)** Schematic of TAT-Cx43_266-283_ treatment and examination at P7. TAT peptide served as control. **(C)** Representative images of TAT or TAT-Cx43_266-283_ treated mouse hippocampus, staining for IBA1 (Scale bar, 100 μm). Plot shows quantification of microglia number in the hippocampus. N = 4 mice. **(D)** Representative images of IBA1+ microglia (Scale bar, 10 μm). Plot shows mean intersection number of microglia (Sholl analysis). N = 44 and 34 cells from 4 mice. **(E)** Representative image of IBA1^+^ microglia, co-staining for FCGR3, C1q, CLEC7A, and MEF2C (Scale bar, 10 μm). **(F)** Plots show quantification of microglial FCGR3, C1q, CLEC7A, and MEF2C intensity in TAT and TAT-Cx43_266-283_ treated mouse hippocampus. N = 4 mice. **(G-H)** Representative images of TAT or TAT-Cx43_266-283_ treated mouse hippocampus, staining for cleaved Caspase 3 or TUNEL labeling (Scale bar, 100 μm). Plot shows quantification of cleaved Caspase 3^+^ or TUNEL^+^ cell number in hippocampus. N = 4 mice. **(I)** Schematic of TAT-Cx43_266-283_ treatment and examination on P14. **(J-K)** Representative fluorescence microscopy images of TAT or TAT-Cx43_266-283_ treated mouse hippocampus, staining for Ki67 or DCX (Scale bar, 50 μm). Plot shows quantification of Ki67^+^ cell number or DCX intensity in hippocampus. N = 4 mice.

**Fig. S6.**
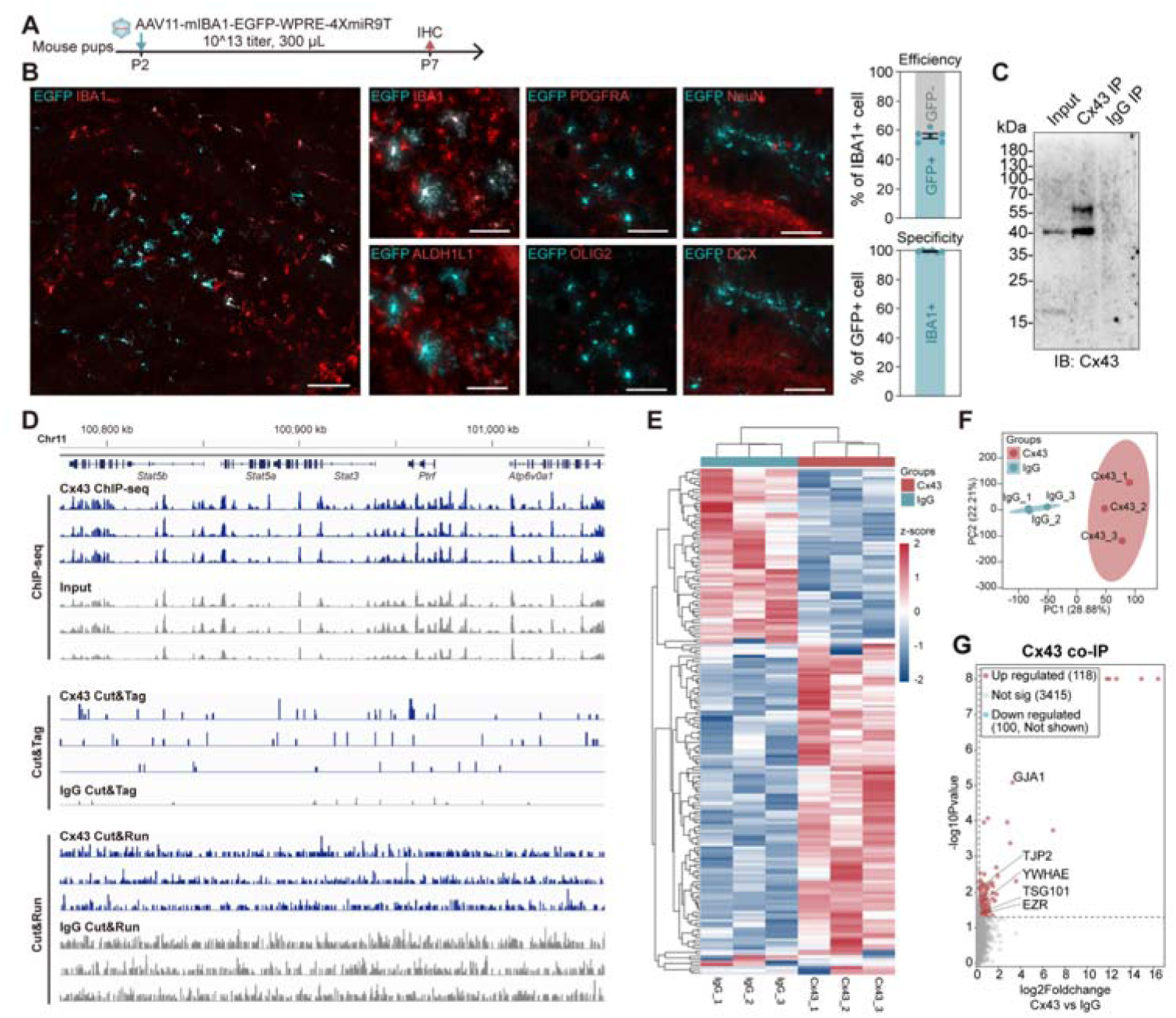
Microglial Cx43 hemichannel did not confer neuroprotective role in development. **(A)** Schematic of AAV-induced Cx43 overexpression experiments. **(B)** Representative images of AAV-transfected mouse hippocampus, staining for GFP and cell markers of microglia (IBA1), astrocyte (ALDH1L1), oligodendroglia (PDGFRA, OLIG2), neuron (NeuN, DCX). (Scale bar, 100 and 50 μm). Plots show quantification of the percentage of microglia transfected by AAV in the peri-injection region (i.e., efficiency), and the percentage of transfected cells that are IBA1+ (i.e., specificity). N = 6 mice. **(C)** Western blot analysis of Cx43 immunoprecipitated product, blotting for Cx43, to verify the IP capacity of antibody. **(D)** Representative Cx43 ChIP-seq, Cut&Tag, and Cut&Run result. **(E)** Heatmap of differentially found proteins in Cx43 co-IP and IgG control. **(F)** PCA analysis of replicates of Cx43 co-IP and IgG control. **(G)** Volcano plot of Cx43 co-IP experiment. Cx43 (GJA1) and its previously reported binding partners were highlighted. Results are expressed as mean ± SEM. \**p* < 0.05, \*\**p* < 0.01.

**Fig. S7.**
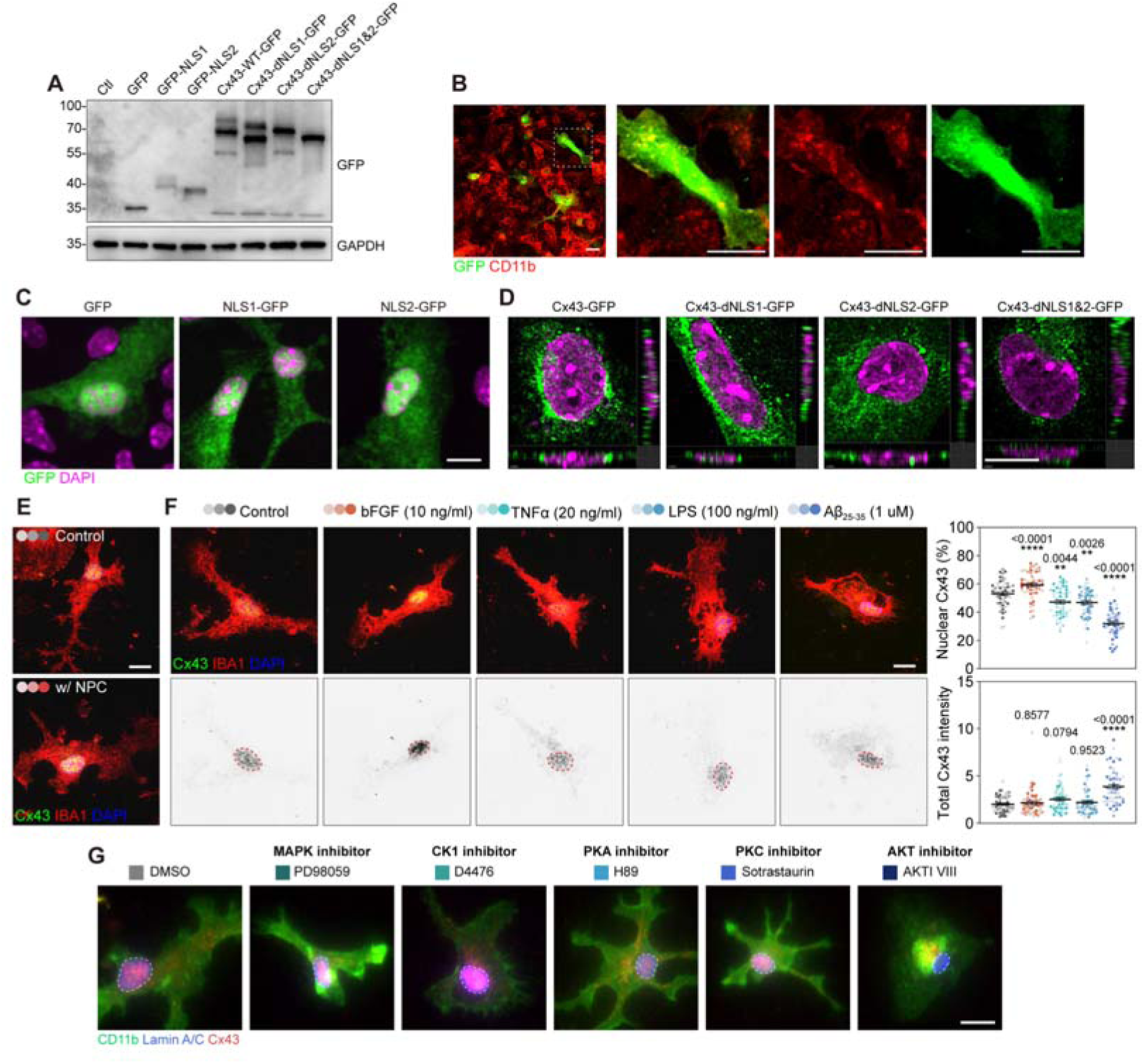
Dynamic nucleus translocation of Cx43 in developmental microglia. **(A)** Western blot analysis of Cx43-NLS-GFP fusion protein overexpression experiment on 293T cell, blotting with GFP antibody, verifying the successful expression of fusion proteins. **(B)** Representative images of lentivirus transfected microglia, staining for CD11b and GFP (Scale bar, 10 μm), verifying successful transfection of microglia. **(C)** Original images of primary microglia in Fig. 7C, with GFP and DAPI staining (Scale bar, 10 μm). **(D)** Original images of primary microglia in Fig. 7D. with GFP and DAPI staining (Scale bar, 10 μm). **(E)** Original fluorescence microscopy images of primary microglia in Fig. 7E, with Cx43, IBA1, and DAPI staining (Scale bar, 10 μm). **(F)** Representative fluorescence microscopy images of primary microglia treated by bFGF, TNFα, LPS, and Aβ (Scale bar, 10 μm). Nuclear Cx43 % and total Cx43 intensity were quantified. Plot shows mean ± SEM. N = 52, 49, 54, 53, 53 cells from 3 experiments. **(G)** Original images of primary microglia in Fig. 7G. with Cx43, CD11b, and Lamin A/C staining (Scale bar, 10 μm). Results are expressed as mean ± SEM. \**p* < 0.05, \*\**p* < 0.01, \*\*\*\**p* < 0.0001.

## Notes

### Competing Interest Statement

The authors have declared no competing interest.

